# Novel features of centriole polarity and cartwheel stacking revealed by cryo-tomography

**DOI:** 10.1101/2020.07.17.208082

**Authors:** Sergey Nazarov, Alexandra Bezler, Georgios N Hatzopoulos, Veronika Nemčíková Villímová, Davide Demurtas, Maeva Le Guennec, Paul Guichard, Pierre Gönczy

**Affiliations:** Swiss Institute for Experimental Cancer Research (ISREC), School of Life Sciences, Swiss Federal Institute of Technology Lausanne (EPFL), Switzerland; Interdisciplinary Centre for Electron Microscopy (CIME), Swiss Federal Institute of Technology Lausanne (EPFL), Switzerland; Department of Cell Biology, University of Geneva, Sciences III, Geneva, Switzerland

**Keywords:** centriole, cartwheel, microtubules, cryo-electron tomography, *Trichonympha*, *Teranympha*, SAS-6

## Abstract

Centrioles are polarized microtubule-based organelles that seed the formation of cilia, and which assemble from a cartwheel containing stacked ring oligomers of SAS-6 proteins. A cryo-tomography map of centrioles from the termite flagellate *Trichonympha* spp. was obtained previously, but higher resolution analysis is likely to reveal novel features. Using sub-tomogram averaging (STA) in *T.* spp. and *Trichonympha agilis*, we delineate the architecture of centriolar microtubules, pinhead and A-C-linker. Moreover, we report ∼25 Å resolution maps of the central cartwheel, revealing notably polarized cartwheel inner densities (CID). Furthermore, STA of centrioles from the distant flagellate *Teranympha mirabilis* uncovers similar cartwheel architecture and a distinct filamentous CID. Fitting the CrSAS-6 crystal structure into the flagellate maps and analyzing cartwheels generated *in vitro* indicates that SAS-6 rings can directly stack onto one another in two alternating configurations: with a slight rotational offset and in register. Overall, improved STA maps in three flagellates enabled us to unravel novel architectural features, including of centriole polarity and cartwheel stacking, thus setting the stage for an accelerated elucidation of underlying assembly mechanisms.

## Introduction

Centrioles are evolutionarily conserved microtubule-based organelles that seed the formation of primary cilia, as well as of motile cilia and flagella. Despite significant progress in recent years, the mechanisms orchestrating centriole assembly remain incompletely understood, in part because the detailed architecture of the organelle has not been fully unraveled.

The centriole is a 9-fold radially symmetric cylindrical organelle typically ∼500 nm in length and ∼250 nm in diameter, which is polarized along a proximal-distal axis (reviewed in Azimzadeh & Marshall, 2010; Gönczy & Hatzopoulos, 2019). In the proximal region lies a likewise symmetrical cartwheel usually ∼100 nm in length, which is critical for scaffolding the onset of centriole assembly (reviewed in Guichard *et al*, 2018; Hirono, 2014). In transverse view, the cartwheel is characterized by a central hub from which emanate 9 spokes that extend towards peripherally located microtubule triplets.

The SAS-6 family of proteins is thought to constitute the principal building block of the cartwheel and is essential for its formation across systems (Culver *et al*, 2009; Dammermann *et al*, 2004; Jerka-Dziadosz *et al*, 2010; Kilburn *et al*, 2007; Kleylein-Sohn *et al*, 2007; Leidel *et al*, 2005; Nakazawa *et al*, 2007; Rodrigues-Martins *et al*, 2007; Strnad *et al*, 2007; Yabe *et al*, 2007). SAS-6 proteins contain an N-terminal globular head domain, followed by a ∼45 nm long coiled-coil and a C-terminal region predicted to be unstructured (Dammermann *et al*, 2004; Kitagawa *et al*, 2011; Leidel *et al*, 2005; Van Breugel *et al*, 2011). *In vitro*, SAS-6 proteins readily homodimerize through their coiled-coil moiety; such homodimers can undergo higher order oligomerization through an interaction between neighboring head domains (Kitagawa *et al*, 2011; Nievergelt *et al*, 2018; Van Breugel *et al*, 2011). Ultimately, this results in the formation of a SAS-6 ring with a central hub harboring 18 juxtaposed head domains, from which emanate 9 paired coiled-coils that extend peripherally. Such ring oligomers are ∼23 nm in diameter and bear striking resemblance with a transverse section of the cartwheel observed in cells. Moreover, recombinant *Chlamydomonas reinhardtii* SAS-6 (CrSAS-6) possesses the ability not only to self-assemble into ring oligomers, but also to undergo stacking of such entities, together generating a structure akin to the cartwheel present in the cellular context (Guichard *et al*, 2017).

Additional features of cartwheel architecture have been unveiled through cryo-electron tomography (cryo-ET) of centrioles purified from *Trichonympha* (Guichard *et al*, 2013, 2012). Three closely related species of these unicellular symbiotic flagellates, *T. campanula, T. collaris* and *T. sphaerica* -referred to collectively as *T.* spp., populate the gut of *Zootermopsis* damp wood termites (Tai *et al*, 2013). *T.* spp. centrioles are particularly well suited for sub-tomogram averaging (STA) of cryo-ET specimens because they harbor an exceptionally long cartwheel-bearing region, reaching several microns (Gibbons & Grimstone, 1960; Guichard *et al*, 2013, 2012). STA of purified *T.* spp. centrioles yielded a ∼34 Å map (Fourier Shell Correlation (FSC) criterion 0.143), which established that the cartwheel comprises stacks of ring-containing elements bearing a central hub from which emanate spokes. Suggestively, a 9-fold symmetrical SAS-6 ring generated computationally from the crystal structure of the CrSAS-6 head domain plus the first 6 heptad repeats of the coiled-coil (CrSAS-6[6HR]) could be fitted in the hub of this STA map (Guichard *et al*, 2012). However, some vertical hub densities remained unaccounted for upon such fitting, raising the possibility that additional components are present. In addition, the *T.* spp. STA map revealed 9-fold symmetrical cartwheel inner densities (CID) inside the hub proper, with contacts between hub and CID occurring where the fitted CrSAS-6[6HR] head domains interact with one another (Guichard *et al*, 2013).

The *T.* spp. STA map uncovered a vertical periodicity of ∼8.5 nm between spoke densities emanating from the hub (Guichard *et al*, 2013, 2012). Two such emanating densities merge with one another as they extend towards the periphery, where the vertical spacing between merged entities is hence of ∼17 nm. There, spokes abut a pinhead structure that bridges the central cartwheel with peripheral microtubule triplets. The STA map also revealed the architecture of the A-C linker, which connects the A-microtubule from a given triplet with the C-microtubule of the adjacent one. Interestingly, both pinhead and A-C linker are polarized along the proximal-distal centriole axis (Guichard *et al*, 2013). Given that the centrally located hub and CID were not noted at the time as being polarized, this led to the suggestion that the pinhead and the A-C linker might be critical for imparting polarity to the entire organelle (Guichard *et al*, 2013). Further cryo-ET analysis of procentrioles from *Chlamydomonas* and mammalian cells established that aspects of A-C linker architecture are evolutionarily conserved, including the attachment points on the A- and C-microtubules; moreover, novel features were revealed, such as a vertical crisscross pattern for the A-C linker in *Chlamydomonas* (Greenan *et al*, 2018; Li *et al*, 2019). Whether the central elements of the cartwheel, including the CID, are likewise conserved beyond *T.* spp. is unclear.

Considering that the earlier work in *T.* spp. was conducted without direct electron detector and that software improvements have occurred since, we sought to achieve a higher resolution map of the *T.* spp. cartwheel-bearing region. Moreover, to explore the evolutionarily conservation of cartwheel architecture, we investigated two other flagellates living in the gut of termites that might likewise harbor long cartwheels well suited for STA.

## Results

### Exceptionally long cartwheel region in *Trichonympha agilis*

We set out to obtain a high-resolution STA map of the native cartwheel in *T.* spp. Moreover, we likewise aimed at investigating *Trichonympha agilis*, a symbiotic flagellate that lives in the gut of the Japanese termite *Reticulitermes speratus* (Ohkuma and Kudo, 1998). This choice was guided by the fact that transcriptomic and genomic information is being assembled in *T. agilis* (Yuichi Hongoh, Tokyo Institute of Technology, Japan, personal communication), which will be instrumental to map proteins onto the STA map should sub-nanometer resolution be reached in the future.

As shown in Fig. 1A, *T. agilis* bears a large number of flagella, which stem from similarly numerous centrioles inserted below the plasma membrane (Kubai, 1973). Many of these flagella are tightly packed in a region called the rostrum located at the cell anterior (Fig. 1A, arrow). To determine the length of the cartwheel-bearing region of *T. agilis* centrioles, cells were resin-embedded and analyzed by transmission electron microscopy (TEM), focusing on the rostral region. Longitudinal views established that the cartwheel-bearing region is ∼2.3 μm in length on average (Fig. 1B, pink line; SD=0.36 μm, N=8). This is less than the ∼4 μm observed in *T.* spp. (Guichard *et al*, 2013), yet over 20 times the size of the ∼100 nm cartwheel in centrioles of most systems, including *Chlamydomonas reinhardtii* and *Homo sapiens* (Guichard *et al*, 2017, 2010; O’Toole and Dutcher, 2014). In addition, we found that the centriole distal region devoid of cartwheel is ∼0.4 μm in *T. agilis* (Fig. 1B, white line; SD=0.07 μm, N=6), similar to its dimensions in other systems (Guichard *et al*, 2013; Le Guennec *et al*, 2020).

**Figure 1:**
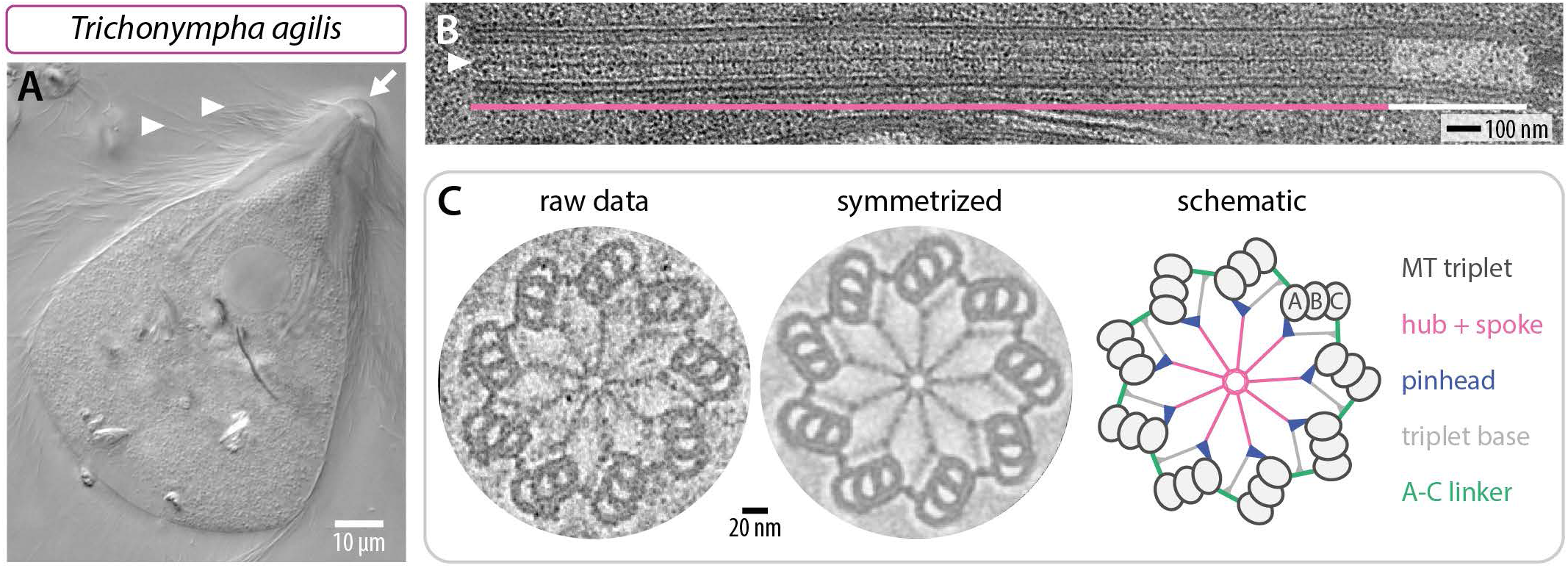
Exceptionally long centriolar cartwheel in *T. agilis*. (A) Differential interference contrast micrograph of live *T. agilis* cell. The arrow points to the cell anterior, where the rostrum is located; arrowheads point to some of the flagella. (B) Transmission electron micrographs of *T. agilis* centriole embedded in resin – longitudinal view; the hub (arrowhead) is visible in the cartwheel-bearing region (pink line), but not in the distal region (white line). (C) Transmission electron micrographs of *T. agilis* centriole embedded in resin in transverse view from distal end (left) and corresponding image circularized and symmetrized with the CentrioleJ plugin (middle), with schematic indicating principal architectural elements (right).

We also analyzed transverse sections of resin-embedded *T. agilis* centrioles using TEM. As shown in Fig. 1C, we found the characteristic features of the cartwheel-bearing region, including a central hub from which emanate 9 spokes that extend towards peripheral microtubule triplets. In addition, we noted the presence of the pinhead and the A-C linker, as well as of the triplet base connecting these two elements (Gibbons and Grimstone, 1960; Vorobjev and Chentsov, 1980), which is more apparent in the circularized and symmetrized image (Fig. 1C).

Overall, given the presence of a long cartwheel-bearing region, we conclude that *T. agilis* also provides a suitable system to investigate the architecture of the proximal part of the centriole using cryo-ET and STA.

### Novel features revealed by improved STA of *Trichonympha* centrioles

Using a direct electron detector, we acquired tilt series of purified *T.* spp. centrioles, focusing on the proximal cartwheel-bearing region (Fig. S1A, S1B), followed by tomogram reconstruction and STA (Fig. 2A-E; for all datasets see Fig. S1C-F for raw tomograms, as well as Fig. S1G-J and Table S1 for processing pipeline). For the central cartwheel, we achieved a local resolution ranging from ∼16 Å to ∼40 Å (Fig. S2A), with a global resolution of ∼24 Å (FSC criterion 0.143; Fig. S2B; see Fig. S3-S5 for resolution of all other STA maps, which have been deposited in EMDB).

**Figure 2:**
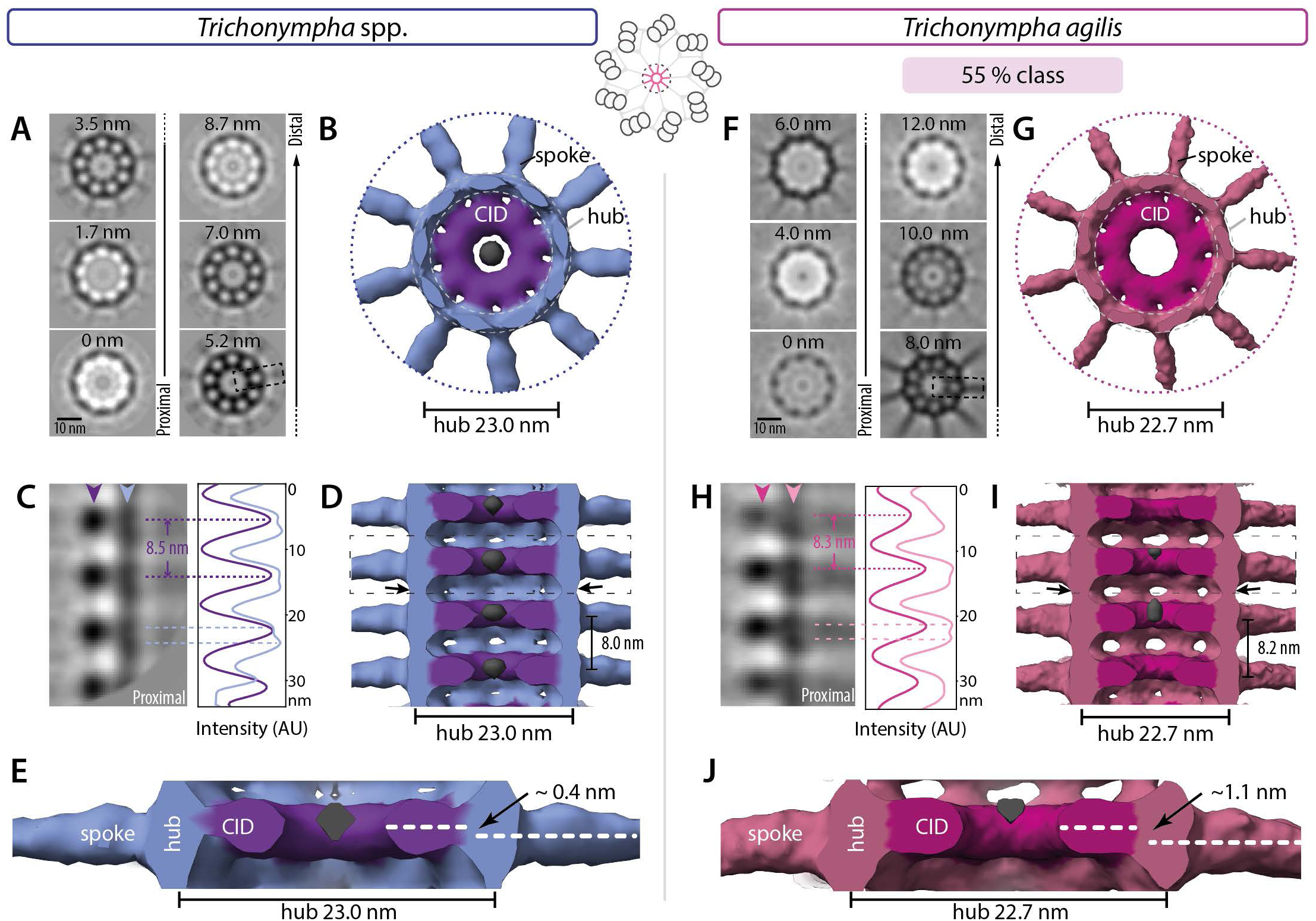
Conserved architecture and polarity in *T.* spp. and *T. agilis* cartwheel. (A, F) Transverse 2D slices through STA of *T.* spp. (A) and *T. agilis* (F) central cartwheel at indicated height, from proximal (0 nm) to distal (8.7 nm (A) and 12 nm (F)). The dashed box in the 5.2 nm (A) and 8.0 nm (F) slices indicate corresponding regions shown in (C, H). Schematic on top illustrates the area used to generate the 3D maps of the central cartwheel. (B, G) Transverse views of central cartwheel STA 3D map; 9 spoke densities emanate from the hub and the CID is present within the hub. The diameter of the hub is 23.0 ± 0 nm in *T.* spp. and 22.7 ± 0.2 in *T. agilis* (both N=3; here and thereafter in the figure legends, N corresponds to 2D measurements from STA averages). An electron dense structure is present inside the CID in *T.* spp. (B, grey), which is also visible to some extent before symmetrizing. Note that the coloring of elements in these and other figure panels is merely illustrative and not meant to reflect underlying molecular boundaries. (C, H) 2D longitudinal view of central cartwheel STA delineated by a dashed box in (A, F) for *T.* spp. (C) and *T. agilis* (H). Arrowheads denote position of line scans along the vertical axis at the level of the CID (C, purple; H, dark pink), and hub (C, light blue; H, light pink), with corresponding pixel intensities in arbitrary units (AU). Plot profiles are shifted horizontally relative to each other for better visibility. Some maxima are highlighted with dashed lines; the average distance between two CID elements is 8.5 ± 0.2 nm (N=4) in *T.* spp. (C) and 8.3 ± 0.5 (N=9) in *T. agilis* (H). Note that hub densities are elongated in the vertical direction, resulting in broad peak profiles where two maxima can sometimes be discerned (dashed light blue lines in (C); dashed light pink lines in (H)). (D, I) Longitudinal views of *T.* spp. (D) and *T. agilis* (I) central cartwheel STA. The average distance between emanating spokes is 8.0 ± 1.5 nm (N=2) in *T.* spp. (D) and 8.2 ± 0.5 nm (N=9) in *T. agilis* (I). Note that these values are measurements on STA and are within the error of the CID periodicities reported in (C, H); the same applies for other measurements hereafter. Discontinuous densities in the center of the CID (grey) are visible in (D, I); the *T. agilis* volume in (I) is shown at a lower contour level than in (G). Boxed area is shown magnified in (E, J). Proximal is down in this and all other figure panels showing STA longitudinal views. (E, J) The CID axis is located distal relative to axis of spoke densities, corresponding to an average shift of 0.9 ± 0.7 (N=3) nm in *T.* spp. (E) and 0.8 ± 0.3 nm (N=9) in *T. agilis* (J).

Using line scans on 2D projections of the STA map, we determined the *T.* spp. hub diameter to be 23 nm (Fig. 2A, 2B), in line with previous work (Guichard *et al*, 2013, 2012). Importantly, the improved resolution achieved here enabled us to uncover novel features in the central cartwheel of the *T.* spp. centriole. Of particular interest, we discovered that the position of the CID is polarized along the proximal-distal centriole axis with respect to the hub and the spokes that emanate from it (Fig. 2C, 2D). This is apparent from vertical intensity profiles of longitudinal views, which show that the CID is positioned distal to the center of the hub density, which itself appears to be elongated in the vertical direction (Fig. 2C). Occasionally, two units can be discerned within one hub density (Fig. 2C, double peaks in light blue intensity profile and corresponding dashed lines), a point that will be considered further below. Such double units were not recognized previously, presumably owing to the lower resolution map (Guichard *et al*, 2013, 2012). Moreover, we found densities that vertically bridge successive hub elements (Fig. 2D, arrows). The polarized location of the CID unveiled here is also apparent with respect to where spoke densities emerge from the hub (Fig. 2E, arrow). In addition, we identified discontinuous densities in the center of the CID (Fig. 2B, 2D).

We likewise analyzed the central cartwheel in *T. agilis* observing variations along the centriole axis in 2D longitudinal views (Fig. S1D). Focused 3D classification of sub-volumes indeed uncovered two classes, corresponding to 55 % and 45 % of sub-volumes, which can occur within the same centriole (Fig. 2F-J; Fig. S6A-F). We found that the central cartwheel STA map of both *T. agilis* classes exhibits many similarities with that of *T.* spp. Thus, the CID is present, and spoke densities emanate from a hub ∼23 nm in diameter (Fig. 2F, 2G; Fig. S6A, S6B). Moreover, vertical densities bridging successive hub elements are also present in both *T. agilis* classes (Fig. 2I, arrows; Fig. S6D, arrows). Furthermore, we found that the CID is also polarized along the proximal-distal centriole axis, being distal with respect to the hub in both *T. agilis* classes, as evidenced from vertical intensity profiles (Fig. 2H-I; Fig. S6C-D), as well as from the location of the CID relative to where spoke densities emerge from the hub (Fig. 2J and Fig. S6D, arrowheads). In the *T. agilis* 55 % class, like in *T.* spp., hub densities are elongated in the vertical direction and can be sometimes discerned as two units (Fig. 2H, double peaks in light pink intensity profile and corresponding dashed lines). The presence of such double hub units next to the CID is more apparent in the 45 % class, where they also exhibit a slight offset relative to the vertical axis (Fig. S6C, white dashed line), a point considered further below. In addition, we found in this class that double hub units alternate with single hub units that do not have a CID in their vicinity (Fig. S6C, dashed arrow, Fig. S6D), an absence noticeable also in raw tomograms (Fig. S1D) and verified in 3D top views (Fig. S6A and S6B, full circle). Moreover, we found that spoke densities emanating from single hub units are thinner than those stemming from double hub units (Fig. S6D). The plausible origin of alternating double and single hub units in the 45 % *T. agilis* class will be considered below. We noted also that the 45% sub-volumes exhibit slight variations in the spacing between double and single hub units (Fig. S6E, 25% and 20% sub-classes).

Taken together, our findings establish that *T.* spp. and *T. agilis* central cartwheel architecture shares many features, including a polarized CID position.

### Comparative STA of peripheral centriole elements in *Trichonympha*

We also investigated peripheral elements in the proximal region of *T.* spp. and *T. agilis* centrioles. To this end, we extracted peripheral sub-volumes from the tomograms and generated for each species three maps using STA centered either on the microtubule triplet, the pinhead or the A-C linker (Fig. 3A, 3E).

**Figure 3:**
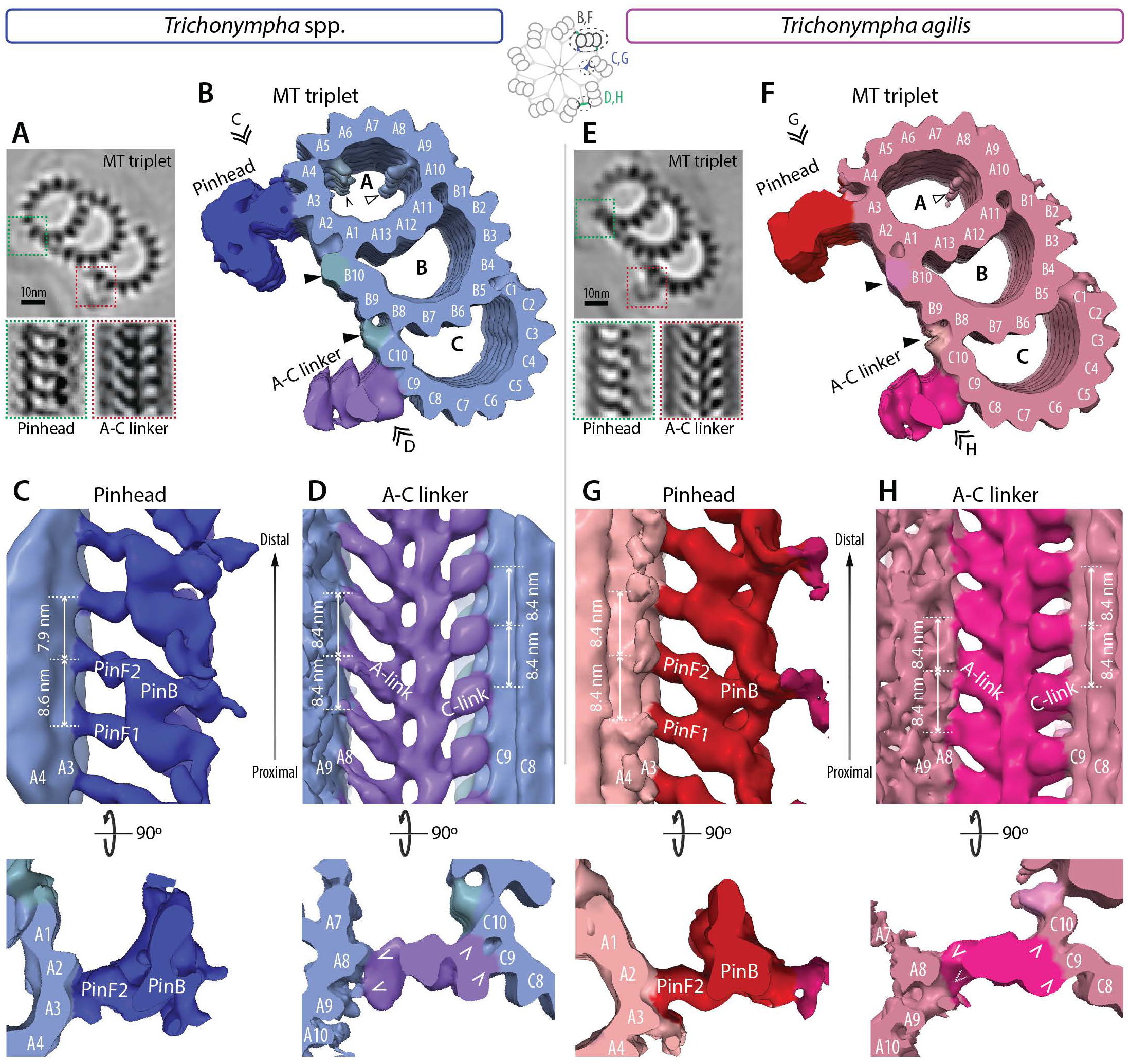
Architecture of peripheral elements in *T.* spp. *and T. agilis.* (A, E) (Top) 2D slices through STA transverse view of microtubule triplet of *T.* spp. (A) and *T. agilis* (E), with insets showing position of pinhead (dashed green box) and A-C linker (dashed red box). (Bottom) Longitudinal 2D slice of STA centered on the pinhead (left) or A-C linker (right). Schematic on top illustrates the different areas used to generate maps of the microtubule triplets (B, F), pinhead (C, G) and A-C linker (D, H). (B, F) Transverse view of *T.* spp. (B) and *T. agilis* (F) microtubule triplet STA. Microtubule protofilament numbers are indicated, as are the pinhead and A-C linker (only the C-link is visible; the A-link lies on the edge of the volume and is thus less well resolved in this STA -for better view see STA centered on A-C linker in D, H). Prominent microtubule inner densities within the A-microtubule are highlighted (empty arrowhead next to A9, chevron next to A5 (B)), as are additional external densities at the A-B and B-C inner junctions (black arrowheads). Double arrowheads point to viewing direction in indicated panels. (C, G) Longitudinal view of STA centered on the pinhead in *T.* spp. (C) and *T. agilis* (G), from the viewing point indicated in (B, F). The pinfeet (PinF1 and PinF2) and pinbody (PinB) are indicated, as are microtubule protofilaments A3 and A4. The average distance between pinfeet elements is 8.6 ± 0.4 nm and 7.9 ± 0.4 nm in *T.* spp. (N=3 each; C) and 8.4 ± 0 nm for each in *T. agilis* (N=3 each; G). Corresponding transverse views are shown below, illustrating the connection of PinF2 with protofilament A3. (D, H) Longitudinal view of STA centered on the A-C linker in *T.* spp. (D) and *T. agilis* (H), from the viewing point indicated in (B, F). Microtubule protofilaments A8/9 and C9/C10 of two adjacent triplets are indicated, as are the A- and C-links. The average distance between A- and C-links is 8.4 ± 0.4 nm and 8.4 ± 0.3 nm in *T.* spp. (both N=6; D) as well as 8.4 ± 0.3 nm and 8.4 ± 0.9 nm in *T. agilis* (N=6 and N=5, respectively; H). Corresponding transverse views are shown below, chevrons point to connecting points; the connection of the A-link with A9 is only partially visible in the transverse view at this height, as indicated by the dashed chevron (H).

For the microtubule triplet, the resulting analysis revealed a characteristic centriolar architecture. Thus, the A-microtubule bears 13 protofilaments, the B-microtubule 10 protofilaments proper, with 3 extra ones shared with the A-microtubule, whereas the C-microtubule exhibits a similar organization as the B-microtubule (Fig. 3B, 3F). Moreover, we detected prominent densities corresponding to microtubule inner proteins (MIPs). In both species, we found a MIP located along protofilament A9, close to protofilament A10 (Fig. 3B, 3F, empty arrowhead). A MIP was discovered at this location in ciliary axonemes (Nicastro *et al*, 2006), and was also observed in centrioles of *T.* spp., *Chlamydomonas* and mammalian cells (Greenan *et al*, 2020, 2018; Guichard *et al*, 2013; Li *et al*, 2019). In addition, we observed a MIP along protofilament A5 in *T.* spp. (Fig. 3B, chevron), which is visible only at low threshold in *T. agilis* and positioned like MIP1 in *Chlamydomonas* centrioles and axonemes (Li *et al*, 2019; Nicastro *et al*, 2006). In both *Trichonympha* species, we also detected additional densities or microtubule associated proteins (MAPs) on the microtubule exterior between the A-B and B-C inner junctions (Fig. 3B, 3F, arrowhead), as described also in *Chlamydomonas* and mammalian centrioles (Greenan *et al*, 2018; Li *et al*, 2019).

For both *T.* spp. and *T. agilis*, we next conducted STA centered on the pinhead or the A-C linker to uncover features in these peripheral elements. We thus found that the pinhead connects to the A3 protofilament in both species (Fig. 3C, 3G), in line with previous observations in *T.* spp. and in other systems (Greenan *et al*, 2018; Guichard *et al*, 2013; Li *et al*, 2019). Moreover, we found that the pinhead is polarized in a similar manner in *T.* spp. and *T. agilis* (Fig. 3C, 3G), with the two pinfeet moieties PinF1 and PinF2 pointing proximally from the A3 protofilament, as reported previously for *T.* spp. (Guichard *et al*, 2013, 2020). We found also that the spacing between PinF1 and PinF2 elements is ∼8.6 nm and ∼7.9 nm in *T.* spp., whereas it is ∼8.4 nm and ∼8.4 nm in *T. agilis* (Fig. 3C, 3G), compatible with power spectra of 2D class averages considering the standard deviation of the measurements (Fig. S2I, S3L).

Similarities between the two species are also apparent in the A-C linker that bridges neighboring MT triplets. Thus, we found that the *T.* spp. A-C linker connects protofilament A8/A9 from one triplet with protofilaments C9/C10 of the adjacent triplet (Fig. 3D), furthering the mapping of these connections compared to previous work (Guichard *et al*, 2013). As shown in Fig. 3H, we found that the A-C linker in *T. agilis* has a similar architecture. To generate an overview of the entire proximal region of the *T. agilis* centriole, comprising hub, spoke, pinhead and A-microtubule, we conducted STA centered on the spokes for both 55% and 45% classes (Fig. S7A-H). Such maps have a lower resolution due to the larger box size, the binning and the absence of radial symmetrization, but nevertheless uncover a concerted proximal-distal polarity of several elements. First, the CID is positioned distally within double hub units. Second, spoke densities exhibit a slightly asymmetry in their tilt angle, with the proximal spoke being more tilted. Moreover, these two polarized features present centrally are connected with the polarized pinhead and A-C linker present peripherally (Fig. S7C-D, S7G-H).

Overall, we conclude that peripheral components of the cartwheel are also generally conserved between the two *Trichonympha* species and exhibit concerted polarity with central elements along the proximal-distal centriole axis.

### Diversity of cartwheel architecture in *Teranympha* cartwheel

The gut of *R. speratus* termites contains another symbiotic flagellate, namely *Teranympha mirabilis* (Koidzumi, 1921; Noda *et al*, 2018), which we found also to harbor numerous centrioles and associated flagella (Fig. S8A). TEM analysis of resin-embedded specimens established that the cartwheel-bearing region in such centrioles is ∼1.1 μm on average (Fig. S8B, green line; N=9, SD=0.06 μm), whereas the distal region averages ∼0.5 μm (Fig. S8B, black line; N=9, SD=0.08 μm). Transverse sections of the cartwheel-bearing region revealed the characteristic hub and spokes connected through a pinhead to peripheral microtubule triplets, which are joined by an A-C linker, whereas the triplet base is poorly visible (Fig. S8C).

We conducted cryo-ET of purified *T. mirabilis* centrioles, acquiring tilt series of the entire cartwheel-bearing region followed by tomogram reconstruction and STA. We again generated separate maps for the central cartwheel, the microtubule triplet, the pinhead and the A-C linker. Focused 3D classification of the central cartwheel region yielded two classes representing 64 % and 36 % of sub-volumes, which can occur within the same centriole (Fig. 4A-G; Fig. S8E). Similarly to the two *Trichonympha* species, a hub ∼23 nm in diameter is present in both *T. mirabilis* classes, with spoke densities emanating from it (Fig. 4A-C). Interestingly, we found that the architecture of the central cartwheel in *T. mirabilis* differs slightly from that in *Trichonympha*. Indeed, in both *T. mirabilis* classes, we uncovered a filamentous structure ∼7 nm in diameter present along the entire cartwheel-bearing region inside the hub, which we dubbed filamentous cartwheel inner density (fCID) (Fig. 4A-G). The fCID is also apparent in transverse views of resin-embedded centrioles and symmetrized tomogram slices (Fig. S8C, S8D). Moreover, the fCID is consistently detected in raw tomograms (Fig. S1E), as well as in non-symmetrized STA comprising larger volumes centered on the spokes (Fig. S7I-P).

**Figure 4:**
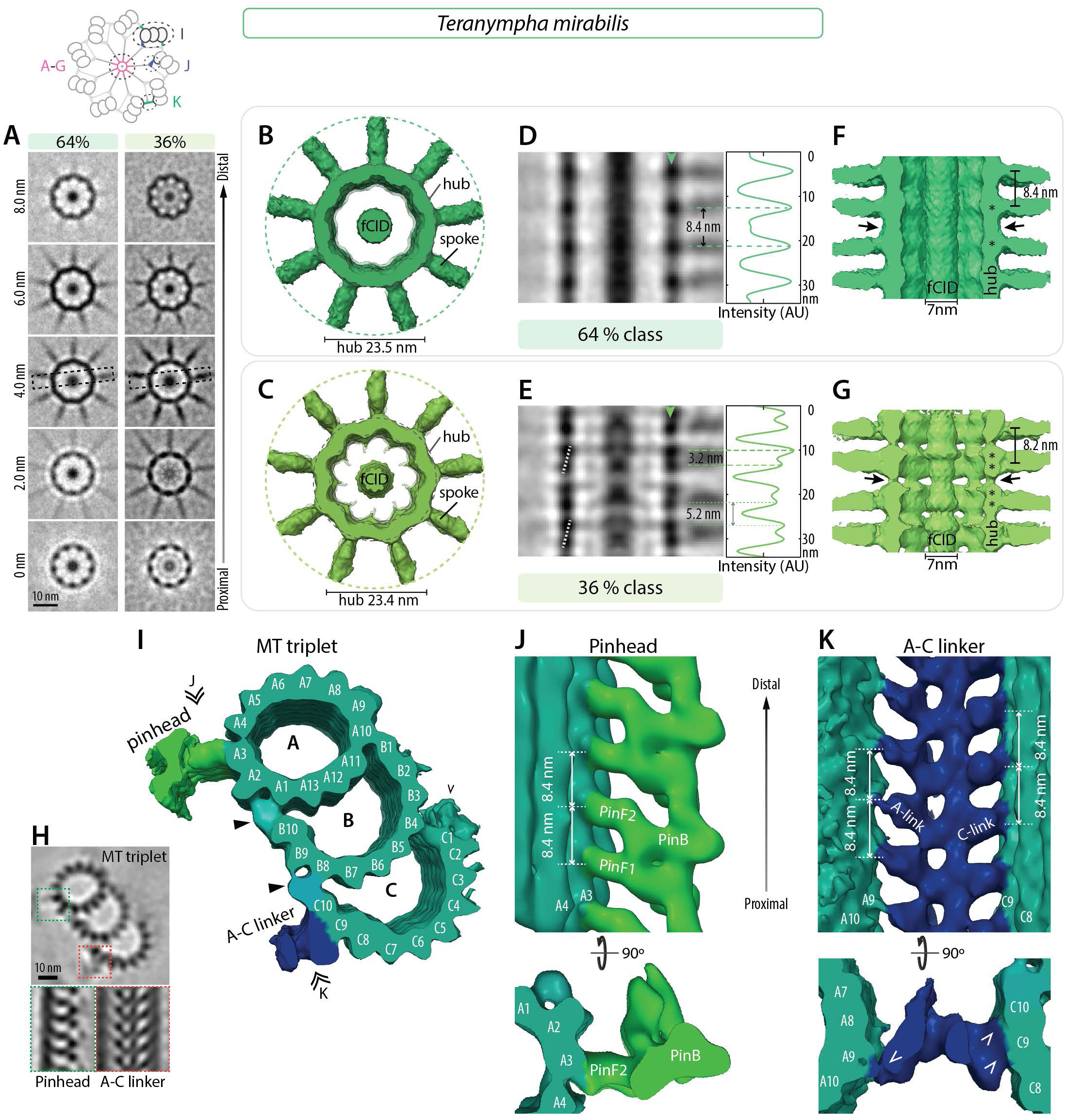
Novel architectural features in *T. mirabilis* cartwheel. (A) Transverse 2D slices through STA comprising 64 % and 36 % of sub-volumes of *T. mirabilis* central cartwheel at indicated height, from proximal (0 nm) to distal (8.0 nm). The dashed boxes in the 4.0 nm slices indicate region shown in D and E. Note filamentous cartwheel inner densities (fCID) in the center of the hub. Schematic on top illustrates the different areas used to generate 3D maps of the central cartwheel (A-G), MT triplets (I), pinhead (J) and A-C linker (K). (B, C) Transverse views of *T. mirabilis* central cartwheel 3D map of 64 % (B) and 36 % (C) classes, with corresponding hub diameters of 23.5 ± 0.2 nm and 23.4 ± 0.5 nm (both N=3). In both classes the fCID is visible and 9 spokes emanate from the hub. (D, E) 2D longitudinal view of *T. mirabilis* central region of the cartwheel STA 64% (D) and 36% (E) classes in the region delineated by dashed boxes in (A). Arrowheads denote position of line scans along vertical axis at the hub level, with corresponding normalized pixel intensities (in arbitrary units). Some maxima are highlighted by dashed lines. The average distance between hub units in the 64 % class is 8.4 ± 0.6 nm (N=8; D); in the 36 % class, the distance between hub densities alternates between 3.2 ± 0.3 nm and 5.2 ± 0.3 nm and (both N=9; E). Dashed white line in (E) indicates the offset between two superimposed hub units at the level of spoke densities, which occurs every other hub unit pair. (F, G) Longitudinal views of *T. mirabilis* cartwheel STA 64 % (F) and 36 % (G) classes. The average distance between spokes is 8.4 ± 0.6 nm (N=8) in the 64 % class and 8.2 ± 1.5 nm (N=7) in the 36 % class. Note in both cases the continuous ∼7 nm fCID inside the hub. Note also densities bridging successive hubs vertically (arrows). Asterisks denote positions of individual units apparent within hub densities. Note that the fCID was not always positioned in the geometrical center of the hub, suggestive of inherent flexibility. (H) (top) 2D slices through STA transverse view of microtubule triplet in *T. mirabilis*, with insets showing pinhead (dashed green box) and A-C linker (dashed red box). (bottom) Longitudinal 2D slice of STA centered on the pinhead (left) or A-C linker (right). (I) Transverse view of microtubule triplet STA in *T. mirabilis*. Microtubule protofilaments are indicated, as are the positions of the pinhead and the C-link (the A-link lies on the edge of the volume and is thus less well resolved in this STA – for better views see STA centered on A-C linker in K). At this contour level, MIPs are not visible. Arrowheads indicate external densities at the A-B and B-C inner junctions; chevron indicates C-stretch. Double arrowheads point to viewing directions shown in (J, K). (J) Longitudinal view of *T. mirabilis* STA centered on the pinhead, from the viewing point indicated in (I). Location of pinhead consisting of pinfeet (PinF1 and PinF2) and pinbody (PinB) are indicated, as are microtubule protofilaments A3 and A4. The average distance between pinfeet elements is 8.4 ± 0.5 nm (N=5). Corresponding transverse view is shown below. (K) Longitudinal view of *T. mirabilis* STA centered on the A-C linker, from the viewing point indicated in (I). Microtubule protofilaments A8/A9 and C9/C10 of two adjacent triplets are indicated, as are the A- and C-links. The average distance between A- and C-links is 8.4 ± 0.9 nm (N=6) and 8.4 ± 0.3 nm, respectively (N=5). Corresponding transverse views are shown below, chevrons point to connections.

As shown in Figure 4D-G, the two central cartwheel *T. mirabilis* classes differ in hub architecture, as revealed by vertical line profile intensity measurements. In the 64 % class, a periodicity of ∼8.4 nm is apparent between hub densities, each consisting seemingly of a single vertically elongated unit (Fig. 4D, 4F). By contrast, in the 36 % class, each hub density comprises a double unit (Fig. 4E, 4G). Such double hub units exhibit a peak-to-peak distance of ∼3.2 nm and are separated from the adjacent double hub unit by ∼5.2 nm (Fig. 4E). The sum of the two distances, namely 8.4 nm is equivalent to that observed in the 64 % class. Moreover, the periodicity at the level of emerging spoke densities is likewise equivalent in the two classes (Fig. 4F, 4G). Furthermore, as in the 45% *T. agilis* class, we observed that every other double hub unit in the 36 % *T. mirabilis* class exhibits a slight offset relative to the vertical axis (Fig. 4E, white dashed lines).

Analysis of the peripheral STA microtubule triplet map of the *T. mirabilis* centriole uncovered the canonical protofilament configuration for A-, B- and C-microtubules (Fig. 4H, 4I). As in *Trichonympha*, we detected additional densities on the external side of the microtubules between A-B and B-C inner junctions (Fig. 4I, arrowheads). The previously described C-stretch that extends from protofilament C1 in *T.* spp. (Guichard *et al*, 2013) is also observed in *T. mirabilis* (Fig. 4I, chevron), while prominent MIPs are not detected at the selected threshold.

The map generated by centering on the pinhead revealed a connection with the A3 protofilament, with PinF1 and PinF2 being separated by 8.4 nm (Fig. 4J), as suggested also by power spectra of 2D class averages (Fig. S4L). Moreover, we found that the *T. mirabilis* A-C linker architecture is similar to that in the two *Trichonympha* species, with anchoring to microtubules at protofilaments A9 and C9/C10 (Fig. 4K).

Overall, these findings indicate that *T. mirabilis* peripheral elements share many conserved features with those in *Trichonympha*, as does the central cartwheel, apart from the discontinuous CID being replaced by the seemingly continuous fCID in *T. mirabilis*.

### Comparative analysis of SAS-6 ring stacking mode

We set out to investigate how SAS-6 rings may fit into the central cartwheel STA maps of the three species. Genomic or transcriptomic information is not available for *T.* spp. and *T. mirabilis* at present. Thus, in the absence of molecular information for SAS-6 in *T.* spp. and *T. mirabilis*, as well as of structural information for *T. agilis* SAS-6, and given that SAS-6 crystal structures from a wide range of organisms exhibit extensive similarities, we employed CrSAS-6 as a model instead. We used homodimers of CrSAS-6[6HR], the longest CrSAS-6 fragment with a determined crystal structure, which can self-assemble into 9-fold symmetrical rings ∼23 nm in diameter and 4.2 nm in height (Kitagawa *et al*, 2011).

Given the presence of two units per hub density in the *T. mirabilis* 36 % class, and also probably in both *Trichonympha* species, we computationally assembled CrSAS-6[6HR] into single rings, as well as into directly superimposed double rings in register. We then used rigid body fitting to score the goodness of fit of both types of ring assemblies with the STA maps (Table S2). Starting with the *T. mirabilis* 36 % class, we found that whereas a single CrSAS-6[6HR] ring fits in this map, additional hub densities remain unaccounted for in this case (Fig. 5A, arrowheads). By contrast, the double ring configuration fills the hub density to a larger extent (Fig. 5B), yielding a better overlap score (Table S2). However, coiled-coil moieties extend slightly outside the spoke densities in this case, potentially because of coiled-coil bending *in vivo* or species-specific features. Moreover, rigid body fitting of CrSAS-6[6HR] homodimers confirmed that two directly superimposed dimers provide a better fit for the *T. mirabilis* 36 % class, a fit that also revealed a slight offset between them (Fig. S9A). Next, we fitted computationally assembled single and double CrSAS-6[6HR] rings in the *T. mirabilis* 64 % class map, finding again that head domains of double rings can be readily accommodated in the hub density, although one coiled-coil moiety clearly extends outside spoke densities in this case (Fig. S9B) Taken together, these observations are compatible with the possibility that SAS-6 rings can directly stack on top of one another *in vivo*.

**Figure 5:**
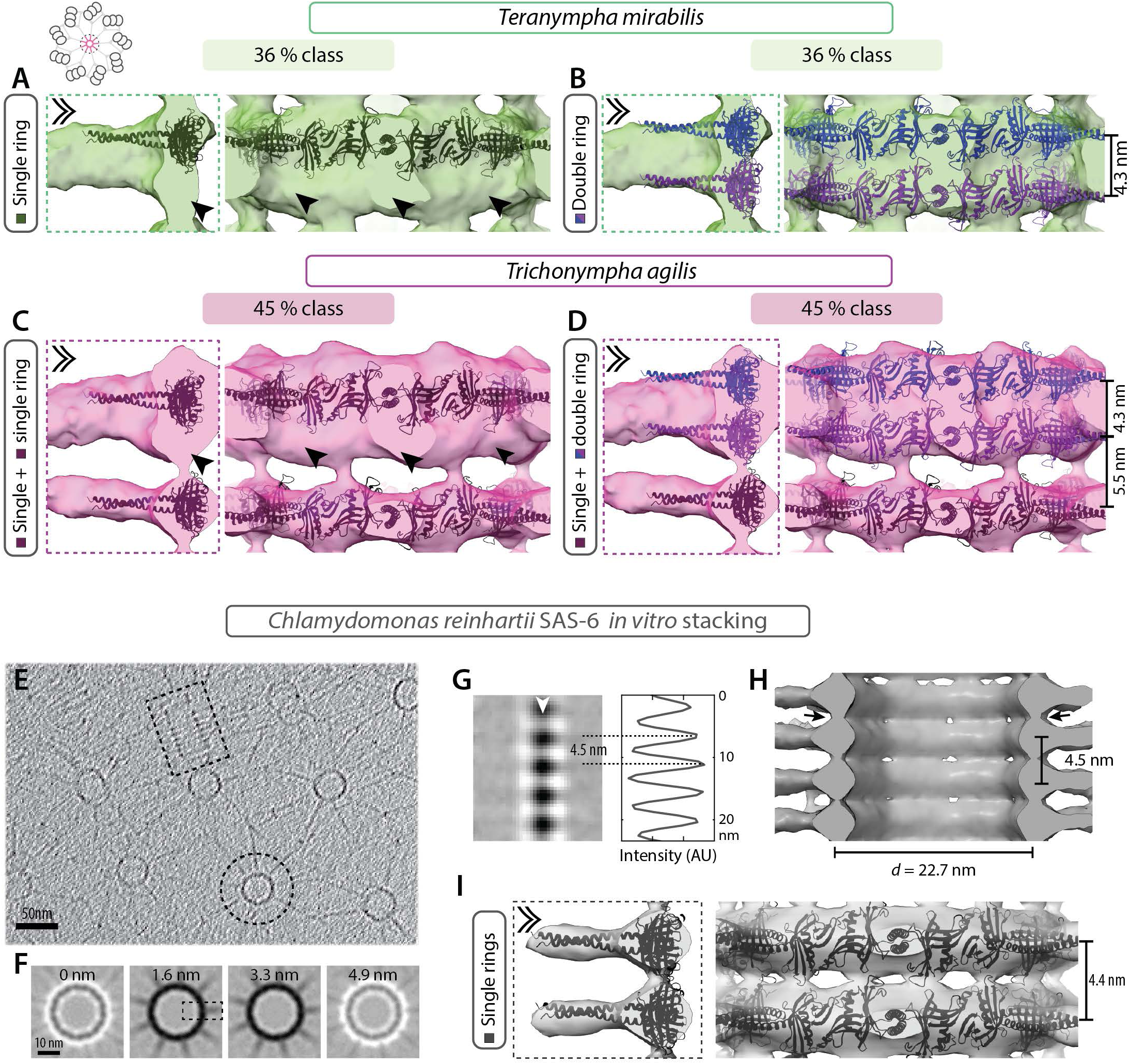
Direct stacking of SAS-6 rings. (A-D) Computationally assembled CrSAS-6[6HR] single (A, C) or double ring in register (ribbon diagram shown in different shades for clarity) (B, D) fitted into the 3D maps of the 36 % *T. mirabilis* class (A, B) and the 45 % *T. agilis* class (C, D). Dashed box shows longitudinal section through hub element (left), double chevron viewing point of longitudinal external views (right). Extra unaccounted densities in single ring fitting are indicated by arrowheads (A, C). Note that each elongated hub density can accommodate two tightly stacked CrSAS-6[6HR] rings (B, D), and that only one ring can fit into the thin hub element in the 45% *T. agilis* class (C, D). Indicated distances stem from measurements on the fitted models. (E) Purified CrSAS-6[NL] protein self-organized into stacks and analyzed by cryo-ET; one longitudinal and one transverse view are highlighted by a dashed rectangle and a dashed circle, respectively. (F, G) 2D views through STA of *in vitro* self-assembled CrSAS-6[NL] proteins. Transverse sections at the indicated heights through one assembly unit (F), as well as longitudinal view (G) of the area delineated by a dashed box in (F); white arrowhead denotes position of line scan along the vertical axis at the level of the hub, with corresponding pixel intensities in arbitrary units (AU). The average distance between two hub elements is 4.5 ± 0.3 nm (N=7; dashed lines). (H) Longitudinal view of *in vitro* self-assembled CrSAS-6[NL] proteins STA 3D map. Note densities bridging successive hubs vertically (arrows). (I) Two CrSAS-6[HR] single rings fitted into the 3D map of *in vitro* self-assembled CrSAS-6[NL] proteins. Dashed box indicates longitudinal section through hub element (left), double chevron indicates viewing point of longitudinal external view (right). Note that the 3D map can accommodate rings stacked 4.4 nm apart (measured on fitted ring models), and vertical densities linking hubs are partially occupied.

To explore whether direct SAS-6 ring stacking might occur also in *Trichonympha*, we likewise performed fitting of computationally assembled CrSAS-6[6HR] single and double rings. We found again that in all cases directly superimposed double rings can readily be accommodated in the hub densities with improved goodness of fit compared to single rings (Fig. 5C, 5D; Fig. S9C, S9D; Table S2). The *T. agilis* 45 % class, which comprises alternating double and single hub elements, presents a particularly interesting case. As anticipated, double rings could be placed in the elongated hub density comprising two units, but a single ring could be accommodated in the thinner individual hub unit (Fig. 5C, 5D; Table S2).

Prompted by the slight offset observed between two SAS-6 ring oligomers upon fitting CrSAS-6[6HR] homodimers in the *T. mirabilis* 36 % class (see Fig. S9A), as well as the structural complementarity between superimposed SAS-6 rings following such an offset, we computationally assembled a double ring with a ∼6.5° rotational offset imposed between ring pairs, which allowed the two rings to come closer to one another vertically by 0.4 nm (Fig. S10B, compare with Fig. S10A; Materials and Methods). Manual fitting in the *T. mirabilis* 36 % class map uncovered that such an offset double ring can be readily accommodated in the hub (Fig. S10C). A similar conclusion was drawn from fitting the offset double ring into the double hub density of the *T. agilis* 45 % class (Fig. S10D). Such a slight offset between superimposed SAS-6 rings might explain the hub offset observed in the longitudinal view of the central cartwheel STA in both *T. mirabilis* 36 % class and *T. agilis* 45 % class (see Fig. 4E and Fig. S6C, white dashed lines). Furthermore, we reasoned that such an offset double ring might likewise affect the connected spoke densities. If spoke densities correspond in reality to two individual spokes slightly offset from one another, as expected from an offset double ring configuration, but which cannot be identified as individual units due to the resolution limit, then the shape of the spoke densities should be elliptical when viewed end on. To investigate this possibility, we unwrapped the central cartwheel maps to obtain a complete view at the level of the spokes (Fig. S10E). Importantly, this uncovered the expected elliptical shape, revealing spoke offset in the *T. mirabilis* 36 % class (Fig. S10F), as well as in both *T. agilis* classes (Fig. S10G) and in *T.* spp. (Fig. S10H). By contrast, no spoke offset was apparent in the *T. mirabilis* 64 % class (Fig. S10F).

Overall, these findings lead us to propose that pairs of SAS-6 rings directly stack in the cellular context and can do so with an offset (see Discussion).

We investigated further the possibility that SAS-6 rings directly stack onto one another using a cell free assay with a recombinant CrSAS-6 protein containing the globular head domain and the entire coiled-coil (referred to as CrSAS-6[NL]). It has been shown previously that CrSAS-6[NL] bearing 6xHis- and S-tag moieties can self-organize into stacks thought to exhibit ∼8.5 nm periodicity between spokes based on the analysis strictly of top views (Guichard *et al*, 2017). We investigated this question anew, now analyzing stacking periodicity of untagged purified CrSAS-6[NL] protein. Moreover, we conducted STA of side views of stacked assemblies, thus allowing us to measure the underlying periodicities with a global resolution of ∼25 Å (Fig. 5E-I, Fig. S5). Importantly, we found that such *in vitro* assembled CrSAS-6[NL] stacks exhibit a periodicity of ∼4.5 nm that can each accommodate one computationally assembled single ring of CrSAS-6[6HR] (Fig. 5G-I). Moreover, we found that the spokes of successive *in vitro* assembled rings are almost in register (Fig. S10I).

Taken together, these observations lead us to propose that direct stacking of SAS-6 rings may be an evolutionarily conserved feature of cartwheel assembly at the onset of centriole biogenesis.

## Discussion

The centriole is fundamental for numerous aspects of cell physiology and shows remarkable conservation across the major eukaryotic groups of life. Understanding the mechanisms that govern centriole assembly requires not only the identification of essential building blocks and their regulators, but also a detailed knowledge of organelle architecture. Our findings uncover conserved features in the cartwheel-bearing portion of the centriole. We find that the CID located within the hub is polarized along the proximal-distal centriole axis in both *Trichonympha* species and that a potentially related fCID is present in *T. mirabilis*. Moreover, we establish that hub densities can be fitted by directly superimposing 9-fold symmetrical SAS-6 double rings that exhibit a slight rotational offset between the two rings. Furthermore, we find that such an offset double ring is stacked in register with the next offset double ring. Overall, our work provides novel insights into the architecture of the centriolar cartwheel at the root of centriole biogenesis.

### Conserved features in cartwheel architecture

Centriole ultrastructure began to be explored in the 1950s when TEM of resin-embedded fixed specimens revealed its signature 9-fold radial symmetry and the chiral arrangement of microtubule triplets (Fawcett & Porter, 1954). More recent cryo-ET analysis uncovered the native architecture of the cartwheel-bearing portion of the centriole (Guichard *et al*, 2013; Li *et al*, 2019). Here, we set out to improve the resolution previously obtained with *T.* spp. and to investigate whether features discovered in this species are present in other flagellates harboring unusually long cartwheels.

In the absence of fossil record or complete sequence information, it is not possible to assess sequence divergence between centriolar proteins in the species analyzed here, nor to date with precision the evolutionary times separating them. However, an approximation is provided in the case of *T.* spp. and *T. agilis* by the phylogenetic divergence between their respective hosts *Zootermopsis* spp. and *Reticulitermes speratus*, which are estimated to have shared their last common ancestor ∼140 million years ago (Bucek *et al*, 2019). Given that *Trichonympha* are obligate symbiotic organisms and considering that the two host species inhabit different continents (Park *et al*, 2006; Thorne *et al*, 1993), it is reasonable to postulate that *T.* spp. and *T. agilis* diverged at least this long ago. *Teranympha mirabilis* belongs to a different genus, and based on small subunit rRNA sequences it is much more distant from the *Trichonympha* species than they are from one another; however, the split between the two genera is thus far undated (Carpenter and Keeling, 2007; Noda *et al*, 2018, 2009; Ohkuma *et al*, 2009). Regardless of the exact evolutionary times separating these flagellates, our findings, together with those of a companion manuscript (Klena *et al*), establish that the CID is conserved between distant eukaryotic groups, suggesting shared evolutionary history and/or function (Fig. 6A). The CID exhibits a 9-fold radial symmetry and connects with the hub approximately where neighboring SAS-6 homodimers interact with one another (Fig. 6B), a suggestive location raising the possibility that the CID imparts or maintains the 9-fold symmetrical SAS-6 ring structure (Guichard *et al*, 2013). We discovered here that the CID is polarized along the proximal-distal centriole axis, so that may also play a role in imparting or maintaining organelle polarity (Fig. 6C). Furthermore, the CID exhibits an ∼8.4 nm periodicity along the proximal-distal centriole axis, with no apparent continuity between two superimposed CID elements. This is in contrast to the fCID, which runs throughout the center of the cartwheel-bearing portion of the *T. mirabilis* centriole. Although the fCID seems disconnected from the hub, small elements linking the two can be discerned in the *T. mirabilis* 36% class (Fig. 6D, see Fig. S7N). Moreover, the fCID might be linked with the hub through other protein segments that are small, flexible or exhibit different periodicities, rendering them difficult to detect in the current STA map. It will be interesting also to address whether the fCID exhibits helical features. Elements related to the fCID may be present in systems other than *T. mirabilis*. Thus, electron-dense material is discernable in the geometrical center of the hub in *T.* spp. and less so in *T. agilis* (see Fig. 2D and 2I), although it is discontinuous and smaller in diameter than the fCID. Moreover, a density that may be related to either CID or fCID is apparent inside the hub in other species, including *Chlamydomonas, Chytrid* and several insects (Geimer & Melkonian, 2004; Guichard *et al*, 2017; O’Toole & Dutcher, 2014; Olson & Fuller, 1968; McNitt 1974; Gottardo *et al*, 2015; Uzbekov *et al*, 2018, Klena *et al*) (Fig. 6A). Determining the molecular nature of CID and fCID in diverse systems is expected to help assess whether they share an evolutionary history and may serve a similar function.

**Figure 6:**
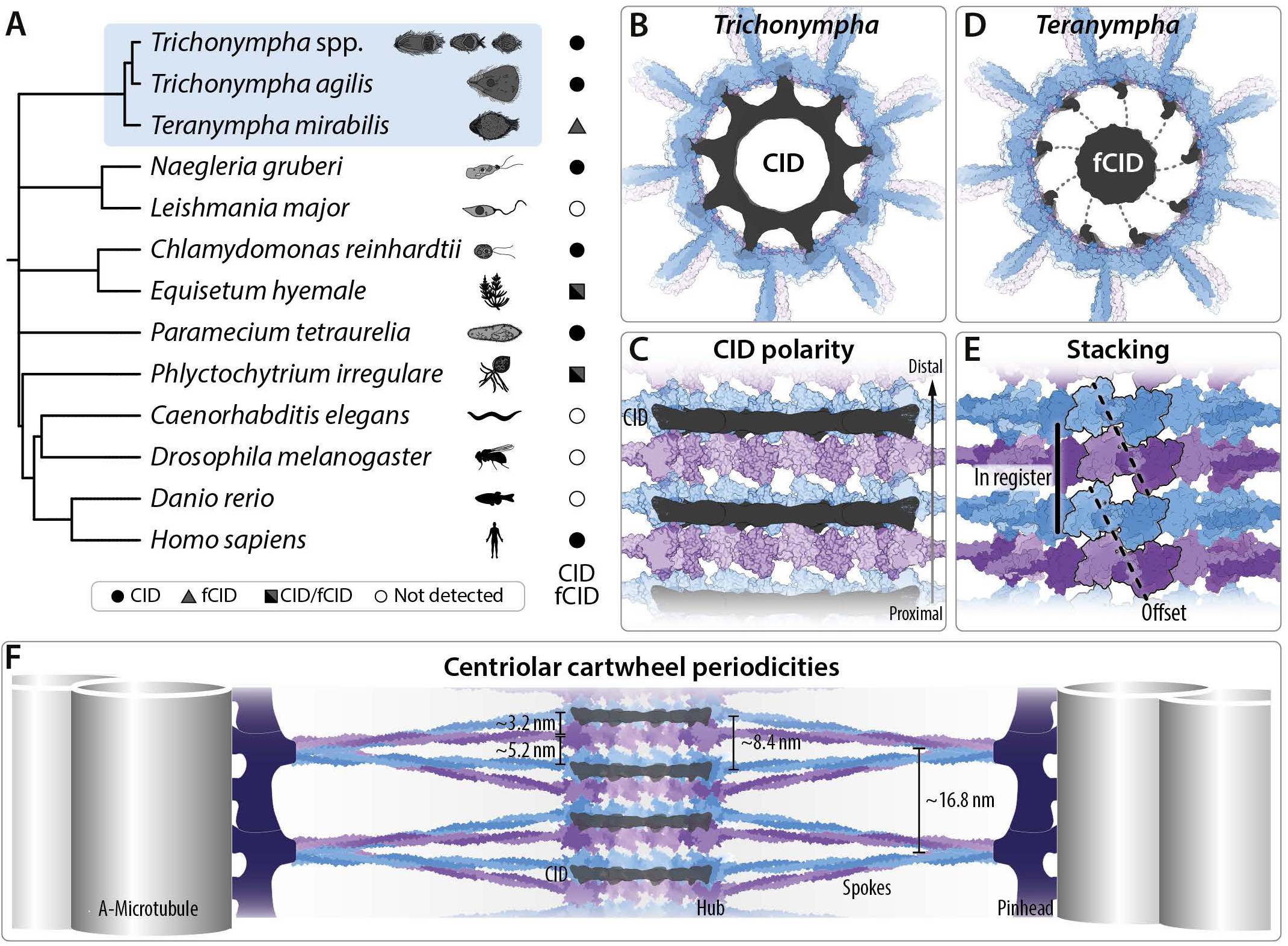
Working model of centriolar cartwheel architecture. (A) Occurrence of CID or fCID in species analyzed in this study (*T.* spp., *T. agilis* and *T. mirabilis*, blue background), and in select other organisms mapped on a taxonomic tree (Tree generated using phyloTv2 based on NCBI taxonomy – https://phylot.biobyte.de/). See main text for references. (B) Schematic of cartwheel hub in *T.* spp. and *T. agilis* with 9-fold symmetrical CID connecting to the hub, equidistant between two spokes. (C) The CID is polarized distally relative to the hub element comprising two tightly superimposed ring elements in *T.* spp. and *T. agilis*. (D) Schematic of cartwheel hub in *T. mirabilis* with continuous fCID along the proximal-distal axis, with potential thin connections to the hub, equidistant between two spokes. (E) Working model of cartwheel stacking, with alternation of two modes of SAS-6 ring stacking: pairs of rings are tightly superimposed with a rotational offset (dashed line) and such offset double rings are then stacked in register (solid line); the offset direction is consistent in *T.* spp., *T. agilis* and *T. mirabilis*, yet may be different in other species, as suggested in *Paramecium* (Klena *et al*). Surface representation of SAS-6 rings; homodimers highlighted by black contour. (F) Consensus model of periodicities in centriolar cartwheel along the proximal-distal axis. Surface representation of offset double rings tightly superimposed with ∼3.2 nm periodicity, in register with the next such offset double ring located ∼5.2 nm away, thus amounting to an overall periodicity of ∼8.4 nm. Note that these values are based on the *T. mirabilis* 36% class, which has the most clearly resolved periodicities, yet does not have a CID as shown in this composite model, with related values in other species; see text for details. The CID exhibits a related ∼8.4 nm mean periodicity in the two *Trichonympha* species. Two spoke periodicities are observed: close to the hub, pairs of spokes are ∼8.4 nm apart, whereas spoke pairs merge towards the periphery, where their periodicity is hence ∼16.8 nm, matching that of the pinhead and connected microtubules. Note that spoke angles are merely illustrative and limited by the resolution of the obtained STA maps. Note also that alternative modes of spoke merging may be revealed by analyzing much larger sub-volumes and be present in other species (Klena *et al*).

We found conserved features also in the peripheral elements of the cartwheel-bearing region of the centriole. Thus, the pinhead is present in all three organisms analyzed here, as is the A-C linker, although the exact arrangement of moieties within these elements differs slightly. Together with the peripheral STA map in the proximal region of the *Chlamydomonas* centriole (Li *et al*, 2019), these findings indicate that peripheral elements have been significantly conserved across evolution. Slight differences between systems are observed notably in the microtubule triplets and prominent MIPs. At the resolution achieved here, we thus identified a prominent MIP in both *Trichonympha* species next to protofilament A9, close to protofilament A10, potentially corresponding to the A-microtubule seam (Ichikawa *et al*, 2017, 2019). A prominent MIP has also been observed at this position in centrioles of mammalian CHO cells and Drosophila S2 cells (Greenan *et al*, 2020, 2018; Li *et al*, 2019). The molecular identity of several *Chlamydomonas* MIPs present in ciliary axonemal microtubules was identified recently (Ma *et al*, 2019), but corresponding MIPs could not be detected here using the selected threshold, perhaps because not all periodicities were explored. Nevertheless, we identified one additional prominent MIP next to protofilament A5 in *T.* spp., which appears to be conserved in metazoan centrioles (Greenan *et al*, 2018), and was identified as RIB72 and FAP252 in ciliary axonemal microtubules (Ma *et al*, 2019). Given the recent spectacular progress in assigning the molecular nature of many ciliary MIPs using high resolution cryo-EM (Ma *et al*, 2019, Song *et al*, 2020), it is likely that better resolution STA maps combined with a search for different periodicities will also enable the efficient identification of centriolar MIPs.

### Root of proximal-distal polarity

The centriole is polarized along its proximal-distal axis, as evidenced for instance by the presence of the cartwheel and microtubule triplets in the proximal region versus microtubule doublets in the distal region. Likewise, distal and sub-distal appendages are restricted to the distal region, by definition. Ultimately, proximal-distal polarity of the centriole also translates into that of the axoneme that emanates from it. Prior work in *T.* spp. raised the possibility that centriole polarity might stem from the pinhead or the A-C linker, because both elements exhibit a clear proximal-distal polarity (Guichard *et al*, 2013). Whereas it remains plausible that polarity is imparted by peripheral elements in some settings, in light of the present findings an alternative view can be envisaged. Indeed, the offset between double SAS-6 rings generates an inherently asymmetric structure, which correlates with the polarized location of the CID, as well as with the asymmetric tilt angle of spoke densities that is also observed in distantly related eukaryotic groups (Klena *et al*). Given that central elements are present before peripheral ones during organelle assembly in the canonical centriole duplication cycle, it is tempting to speculate that the asymmetric properties of offset double rings and/or the CID are key for imparting organelle polarity. The situation might differ during *de novo* centriole assembly, which in human cells does not require interactions between HsSAS-6 head domains (Wang *et al*, 2015). In this case, peripheral elements are thought to be more critical, and this may also be the case when it comes to imparting organelle polarity.

### Working model of SAS-6 ring stacking

What are the mechanisms of SAS-6 ring stacking? The previous lower resolution *T.* spp. map (Guichard *et al*, 2013), together with an initial analysis of *in vitro* generated stacks of CrSAS-6[NL] (Guichard *et al*, 2017), led to the suggestion that the periodicity between subsequent SAS-6 ring oligomers was ∼8.5 nm. The higher resolution analysis performed here in three species leads us to propose instead that SAS-6 rings stack directly on top of one another, thus doubling the number of SAS-6 molecules per vertical unit length. Although we cannot formally exclude that a single SAS-6 ring is present and that additional densities correspond to other unassigned protein(s), several pieces of evidence support the direct SAS-6 stacking scenario. First, two SAS-6 rings fit without clashes within vertically elongated hub densities in *Trichonympha* and *T. mirabilis*. Second, the *T. agilis* 45% class contains thin hub densities that can accommodate only one SAS-6 ring, reinforcing the notion that the thicker ones harbor two. Third, stacks reconstituted in a cell free assay from purified CrSAS-6 exhibit direct ring superimposition.

Our findings taken together lead us to propose the following working model of SAS-6 stacking in the cellular context (Fig. 6E, 6F). This model entails two modes of ring stacking. The first corresponds to the offset double ring configuration, with a tight packing of SAS-6 rings, which are ∼3.2 nm apart for instance in the *T. mirabilis* 36% class. In line with our findings, directly superimposed hub densities with remarkably similar spacing are observed also in *Paramecium* and *C. reinhardtii* (Klena *et al*). In the second stacking mode, two such offset double rings stack in register, being ∼5.2 nm apart from each other in the *T. mirabilis* 36% class. A similar alternating pattern is observed in *T.* spp. and *T. agilis*, albeit with slightly different distances, potentially owing to limitations in measurement precision. Overall, we propose that these two alternating modes of SAS-6 ring stacking represent a fundamental feature of cartwheel assembly.

The periodicity of the basic central cartwheel unit in this working model is ∼16.8 nm, which corresponds to the largest conserved periodicity detected by FFT. This value is consistent with the distance expected from an alternation of the two modes of SAS-6 stacking (2x ∼3.2 nm for the rings with an offset + 2x ∼5.2 nm for the rings in register in the case of the *T. mirabilis* 36% class = ∼16.8 nm) (Fig. 6F). Each SAS-6 hub element possesses one spoke emerging from it, but owing to insufficient resolution, two closely superimposed spokes appear as one joint density in all cases, except for the single hub element in the *T. agilis* 45% class. Interestingly, we note also that hub elements with accompanying spoke densities are further apart in the *T. mirabilis* 64 % class, indicative of a potential variation in stacking mode. Perhaps the densities bridging successive hub elements in this case correspond to another protein than SAS-6. One intriguing candidate to consider in this context is SAS-6-like, an evolutionary ancient SAS-6 related protein lacking the coiled-coil (de Leon *et al*, 2013).

### Concluding remarks

In summary, conducting cryo-ET in three flagellates enabled us to uncover important conservation of centriolar architecture, as well as some species specificity. In the future, higher resolution analyses, including without symmetrizing, is expected to reveal additional polarity and stacking features, as well as to enable one to visualize protein domains and thus further unravel the mechanisms governing centriole assembly.

## Materials and methods

### Transmission electron microscopy of *Trichonympha and Teranympha* cells

*T. agilis* and *T. mirabilis* were extracted from the hindgut of *Reticulitermes speratus* termites (a kind gift of Yuichi Hongoh, Tokyo University of Technology, Japan), as described (Guichard *et al*, 2015), placed in a drop of 10 mM K-PIPES, and gravity-sedimented for 3-5 min onto coverslips coated with poly-D lysine (Sigma-Aldrich, catalog number P1024), until cells became in contact with the glass. Cells were fixed overnight in 2 % paraformaldehyde 1 % glutaraldehyde in phosphate buffer 0.1 M, pH 7.4 (with 0.2 % Tween-20 for some samples), washed in cacodylate buffer (0.1 M, pH 7.4) at 4°C and post-fixed with 0.5 % tannic acid in cacodylate buffer (0.1 M, pH 7.4) for 40 min at room temperature. After two washes in distilled water, cells were post-fixed in 1% osmium tetroxide in cacodylate buffer. Thereafter, samples were washed in distilled water and dehydrated in a graded ethanol series (1× 50 %, 1× 70 %, 2× 96 %, 2× 100 %). Finally, samples were embedded in epon resin and polymerized at 65°C overnight. 50 nm sections were cut and then stained in 1% uranyl acetate in distilled water for 10 min, followed by Reynold’s stain (1.33 g lead citrate, 1.76 g sodium citrate, 160 mM NaOH in 50 ml distilled water (Reynolds, 1963)) for 5 min. Images were acquired using Tecnai Spirit at 80 kV or Tecnai F20 at 200 kV microscopes (Thermo Fischer Scientific). Pear-shaped *T. agilis* cells with long flagella covering the anterior of the cell can be readily distinguished from elongated *T. mirabilis* cells with shorter flagella arranged in rows spiraling around the cell. These distinct morphologies are preserved in resin. Purified centrioles from the two species were distinguished in cryo-ET based on the presence or absence of the fCID. Images of centriole cross-sections from TEM and cryo-ET were circularized and symmetrized using the CentrioleJ plugin (Guichard *et al*, 2013).

### Purification of *Trichonympha and Teranympha* centrioles

*Trichonympha* and *Teranympha* cells were extracted from the hindgut of termites in 10 mM K-PIPES in the presence of cOmplete protease inhibitor cocktail (1:1000; Roche) as described (Guichard *et al*, 2015), with minor species-specific modifications. For the purification of centrioles from mixed *T. agilis* and *T. mirabilis* populations from *R. speratus* termites, cells were pelleted at 1000 g for 20 s and the supernatant discarded. For enrichment of *T. mirabilis* centrioles, cells were sedimented on ice 3×10 min in 1 ml 10 mM K-PIPES; the more fragile *T. agilis* cells spontaneously lysed during this step and thus depleted from the resulting pellet. *T.* spp. (*T. collaris, T. campanula* and *T. sphaerica*) were extracted from *Zootermopsis angusticollis* termites (a kind gift from Filip Husnik and Patrick Keeling, University of British Columbia, Vancouver, Canada), which harbor the same set of *Trichonympha* species as *Zootermopsis nevadensis* (Boscaro *et al*, 2017; Tai *et al*, 2013). After extraction, *T.* spp. cells were sedimented on ice 3×10 min in 1 ml 10 mM K-PIPES. In all cases, centrioles were released during a 20-40 min incubation on ice in 1 ml K-PIPES 10 mM 0.5 % NP-40 plus protease inhibitors. Centrioles and associated flagella were either pelleted at 500 g for 3 min or the resulting supernatant spun at 1000 g for 5 min at 4° C; both types of preparations were utilized for cryo-ET. In all cases, centrioles were washed once in 1 ml 10 mM K-PIPES plus protease inhibitors and pelleted at 1000 g for 5 min. The pellet was stored on ice before preparing grids for cryo-ET.

### Cryo-ET data acquisition

Grids were prepared as described previously (Guichard *et al*, 2015). Briefly, 4 μl of the purified centriole solution were mixed with 10 nm gold beads using a 20 μl tip cut at the extremity and then applied to the copper side of a glow discharged lacey holey carbon grid (carbon Cu 300 mesh, Ted Pella). The grid was then manually blotted from the carbon side with a Whatman grade 1 filter paper and plunged into liquid ethane at −180° C with a manual plunge freezer.

Tilt series of centrioles that were approximately parallel to the tilt axis were collected with the Tomo 4.0 software (ThermoFisher Scientific) on a Tecnai F20 TEM operated at 200 kV (ThermoFisher Scientific); data were recorded with a Falcon III DD camera (ThermoFisher Scientific) in linear mode at 29’000X magnification, corresponding to a pixel size of 3.49 Å. The tilt series were recorded from −60° to 60° using both continuous and bi-directional tilt schemes, with an increment of 2° at 2.5–5 µm underfocus. To allow determination of the proximal-distal axis, the blunt proximal end was included in image acquisition. A total of 54 tilt series were collected (Table S1).

Tilt series alignments using gold fiducials were performed with IMOD v4.9 (Kremer *et al*, 1996). The contrast transfer function (CTF) was estimated with CTFFIND4 v1.8 (Rohou and Grigorieff, 2015) using relion_prepare_subtomograms.py (Bharat and Scheres, 2016). Variable SkipCTFCorrection was set to ‘True*’* to generate a wedge-shaped tilt- and dose-weighted 3D CTF model. Aligned tilt series were corrected by phase-flipping with ctfphaseflip and used for weighted back-projection (WBP) tomogram reconstruction with IMOD.

### Sub-tomogram processing

Areas corresponding to the central cartwheel region (CW) and to the peripheral microtubule triplet region (MTT) were identified visually in individual tomograms and their longitudinal axes modeled with closed contours in 3dmod. Proximal-distal polarity of the centriole was assessed by the fact that the proximal end was blunt, in contrast to the distal end, where the flagellum was usually present. Moreover, the clockwise chirality of the microtubule triplets when viewed from the distal end provided an independent means to determine proximal-distal polarity, which was registered and maintained in the resulting sub-volumes. Also, reconstructions from sub-volumes re-centered on the spokes with CW and polar CID density on one side, and spokes and polar pinhead on the other, allowed unambiguous polarity assignment (see Fig. S7).

Individual model points were interpolated along the contours using the addModPts from the PEET package (Heumann *et al*, 2011), with a step size of 3 × 85 Å = 252 Å corresponding to a 73 pixels shift (see Fig. S1B). The interpolation step size was selected to correspond to approximately three hub element, as smaller step sizes could not separate cartwheel variations visible in the raw tomograms (see Fig. S1D). Larger step sizes were also explored to generate non-overlapping sub-volumes along the proximal-distal axis of the cartwheel hub, shifted relative to each other by more than half of the sub-volume size. The resulting STAs exhibited similar structural features, yet were noisier due to the reduced number of sub-volumes.

Interpolation with a step size of 252 Å resulted in 1385 initial CW sub-volumes for *T.* spp., 1154 for *T. agilis* and 1802 for *T. mirabilis*. A similar approach and step size was selected for extraction of MTT, resulting in 2792 initial MTT sub-volumes for *T.* spp., 3850 for *T. agilis* and 7928 for *T. mirabilis* (see Fig. S1G-I; Table S1). All further processing was performed with RELION v2.1 or v3.1 (Kimanius *et al*, 2016). Briefly, 2D projections of all sub-volumes along the Z axis were generated and subjected to reference-free 2D classification. 2D class averages with clear CW or MTT densities were selected for further data processing. At this step, the hub of both *T. agilis* and *T. mirabilis* showed variations in vertical spoke and hub element spacing. Power spectra of 2D class averages were used to measure the vertical spacing of the CW elements (see Fig. S2C, S3C, S3F, S4C, S4F) or MTT (see Fig. S2F, S3I, S7I). Modeling with 3dmod was also used to investigate the proximal-distal twist of microtubule blades. No twist angle of the microtubules was observed along the proximal-distal axis of the cartwheel-bearing region in either species, as previously reported for *T.* spp. (Gibbons and Grimstone, 1960).

Selected 2D projections of CW sub-volumes were re-extracted, now as 3D sub-volumes with a box size of 200 pixels (corresponding to 80 nm, with the exception of *T.* spp., which corresponds to 88 nm) and re-scaled to a pixel size of 4 Å. Initial 3D CW and MTT references were built *de novo* with the Stochastic Gradient Descent (SGD) algorithm from the re-extracted 3D sub-volumes. The resulting CW initial reference was re-aligned with relion_align_symmetry to orient the visible nine-fold rotational (C9) axis along Z axis, and C9 symmetry applied. Similarly, the MTT initial reference proximal-distal axis was re-aligned with the Z axis. Consensus 3D refinement of the CW initial reference with C9 symmetry and missing wedge correction revealed a well resolved CW density. Consensus 3D refinement was followed by focused 3D refinement with a soft mask that included only the CW hub with the CID, as well as part of the emanating spoke density, whose inclusion inside the mask was important to avoid misalignment along the proximal-distal axis. Moreover, one round of focused 3D classification in *T. agilis* and *T. mirabilis* with a soft tight mask that included only three central stacked rings revealed vertical spacing variations in 3D.

The *T. agilis* CW hub classified into two groups representing 55 % and 45 % of refined sub-volumes. The 55 % class showed uniform 85 Å vertical spacing measured from the 3D reconstruction in real space and from the power spectrum. Further 3D classification did not yield any observable improvement in the quality of the maps as judged by features and resolution. In contrast, further focused 3D classification without alignment of the 45 % class revealed two sub-groups corresponding to 20 % and 25 % of all refined sub-volumes, with differences in distances between alternating thin and thick rings. The *T. mirabilis* CW hub classified into two main groups representing 64 % and 36 % of refined sub-volumes. The 64 % class exhibited 85 Å vertical spacing measured from the 3D reconstruction in real space and from power spectra and a smooth CW hub surface. In contrast, the 36 % class revealed 85 Å vertical spacing with two smaller units per hub with 42 Å spacing measured from the 3D reconstruction in real space and from power spectrum. Further 3D classification did not yield any observable improvement in the quality of the maps as judged by features and resolution. For all 3D classes, final focused 3D refinements with local alignments were performed. Analogous analyses of *T.* spp. revealed a single stable class with uniform 85 Å vertical spacing measured from the 3D reconstruction and from the power spectrum.

To establish the polar relationship between the CID, the corresponding CW and the peripheral elements (A-microtubule, pinhead, A-C linker), the symmetry was relaxed from C9 to C1 with relion_particle_symmetry_expand. The resulting symmetry-expanded sub-volumes were re-extracted with binning 3 and a box size of 100 pixels, using the ‘re-center refined coordinates’ option approximately on the spokes center to include CW density on one side and spokes and pinhead on the other. Focused 3D classification of re-extracted sectors without alignment and without symmetry iterated with focused 3D refinement revealed reconstructions with well-resolved connections between central and peripheral elements. No structural features with a periodicity different than a multiple of 85 Å were detected at this step.

A similar processing pipeline was applied to peripheral elements (microtubules, pinhead, A-C linker). Selected 2D projections of MTT sub-volumes were re-extracted as binned 2 times 3D sub-volumes with a box size of 128 pixels corresponding to 89 nm to include microtubule triplet and emanating pinhead and A-C linker densities (see Fig. S1B). Initial 3D MTT references were built *de novo* with the SGD algorithm from the re-extracted 3D sub-volumes. The resulting MTT initial reference proximal-distal axis was re-aligned with the Z axis. Re-extracted MTT sub-volumes were subjected to consensus 3D refinement followed by rounds of focused 3D classification and 3D refinement. Refined sub-volumes were re-extracted either at the center of emanating pinhead or A-C linker densities and refined separately. Local resolution distributions were determined with ResMap (Kucukelbir *et al*, 2014) and directional 3D FSC were measured with 3DFSC web server (Zi Tan *et al*, 2017).

### Rigid body fitting and computational assembly of SAS-6 rings

Rigid-body fitting of CrSAS-6[6HR] (PDB-3Q0X) into *T.* spp., *T. agilis and T. mirabilis* 3D STA maps was performed with ADP_EM plugin (Garzón *et al*, 2007), followed by symmetrical fitting in UCSF Chimera (Pettersen *et al*, 2004).

Single CrSAS-6[6HR] rings were computationally assembled as described (Kitagawa *et al*, 2011), while double rings were assembled by superposing two such CrSAS-6[6HR] rings using the crystallized stacks of LmSAS-6 rings as guide (Van Breugel *et al*, 2011). Offset double rings were assembled manually by rotating the proximal ring clockwise, with the angle being estimated based on angles between fitted homodimers. The vertical distance between rings in the double offset ring configuration was reduced by 0.4 nm, and clashes were only allowed between flexible loops that are not resolved in the crystal structure. CrSAS-6[6HR] single ring, double in register rings or double offset rings were fit in the map using *fitmap* function of ChimeraX (Goddard *et al*, 2018). Coordinate models were initially converted to molecular maps at corresponding global resolutions and then fitted into the STA maps. Calculated values for correlation and overlap between maps are reported in Table S2. UCSF Chimera or ChimeraX were used for visualization and segmentation of 3D models.

### *In vitro* assembly of CrSAS-6 stacked cartwheels

A 6xHis- and S-tag-containing CrSAS-6 construct harboring the N-terminal globular head domain plus the entire coiled-coil (referred to a CrSAS-6[NL], comprising amino acids 1-503) was expressed and purified as described (Guichard *et al*, 2017), except that the tag moieties were removed by TEV protease cleavage. The resulting purified CrSAS-6[NL] protein was dialyzed overnight from Tris pH 7.5 150 mM NaCl into 10 mM K-PIPES pH 7.2 using a mini dialysis unit at 4°C (slide-A-lyzer, 3.5 K, 10–100 ml, Pierce, catalogue number 69’550). Thereafter, 5 μl of dialyzed material was pipetted onto a glow-discharged Lacey holey carbon grid (Carbon Cu 300 mesh, Ted Pella), blotted for 3 s (blot force −15, no wait time) and plunge-frozen by vitrification in liquid nitrogen-cooled liquid ethane, using a Vitrobot MKIV (ThermoFisher Scientific).

Tilt series were collected on a Titan Krios TEM (ThermoFisher Scientific) operated at 300 kV and equipped with a Gatan Quantum LS energy filter (zero loss slit width 20 eV; Gatan Inc.) on a K2 Summit direct electron detector (Gatan Inc.) in electron counting mode (110 e^−^/Å^2^ total dose), at a calibrated pixel size of 2.71 Å. The tilt series were recorded from −60° to 60° using a dose-symmetric tilt scheme (Hagen *et al*, 2017) with increments of 2° at 2.5-3.5 µm defocus. A total of 31 tilt series were collected automatically using SerialEM (Mastronarde, 2005).

Tilt series alignment using gold fiducials was performed with IMOD v4.9. The CTF was estimated with CTFFIND4 v1.8 using relion_prepare_subtomograms.py. Variable SkipCTFCorrection was set to ‘True’ to generate a wedge-shaped tilt- and dose-weighted 3D CTF model. Aligned tilt series were corrected by phase-flipping with ctfphaseflip and used for WBP tomograms reconstruction with IMOD. Further STA was performed as described above for *Trichonympha* centrioles. Briefly, side views of CrSAS-6[NL] stacks were identified visually in tomograms and their longitudinal axes modeled with closed contours in 3dmod. Next, individual model points were added along the contours using addModPts with a step size of 2 CrSAS-6[NL] rings (2 × 42 Å = 84 Å) to prevent misalignment of potentially merged spokes, resulting in 926 initial sub-volumes (see Fig. S1J; Table S1). Further processing was performed with RELION v2.1 or v3.1 as described above.

## Declarations

### Ethics approval and consent to participate

Not applicable.

### Consent for publication

Not applicable.

### Availability of data and material

Sub-tomogram averages have been deposited at Electron Microscopy Data Bank (EMD-10916, EMD-10918, EMD-10922, EMD-10923, EMD-10927, EMD-10928, EMD-10931, EMD-10932, EMD-10934, EMD-10935, EMD-10937, EMD-10938, EMD-10939, EMD-10941, EMD-10942; Table S1).

### Authors’ contributions

SN, AB, GNH, VNV and DD collected and analyzed data. SN, AB, GNH, PG contributed to experimental design and analysis, as well as writing the manuscript. PaG and MLG contributed to data analysis. All authors read and approved the final manuscript.

## Acknowledgements

We are grateful to Yuichi Hongoh, Patrick Keeling and Filip Husnik for providing termites. We thank Graham Knott and Stéphanie Rosset (BioEM platform of the School of Life Sciences, EPFL) for assistance with TEM, as well as Kenneth Goddie, Lubomir Kovacik and Henning Stahlberg (Center of Nanoimaging, Biozentrum, Basel, Switzerland) with data collection on the Titan Krios. Niccolò Banterle, Graham Knott and Fabian Schneider are acknowledged for their critical reading of the manuscript.

## Additional Data

### Supplemental Table Legends

**Table S1.**
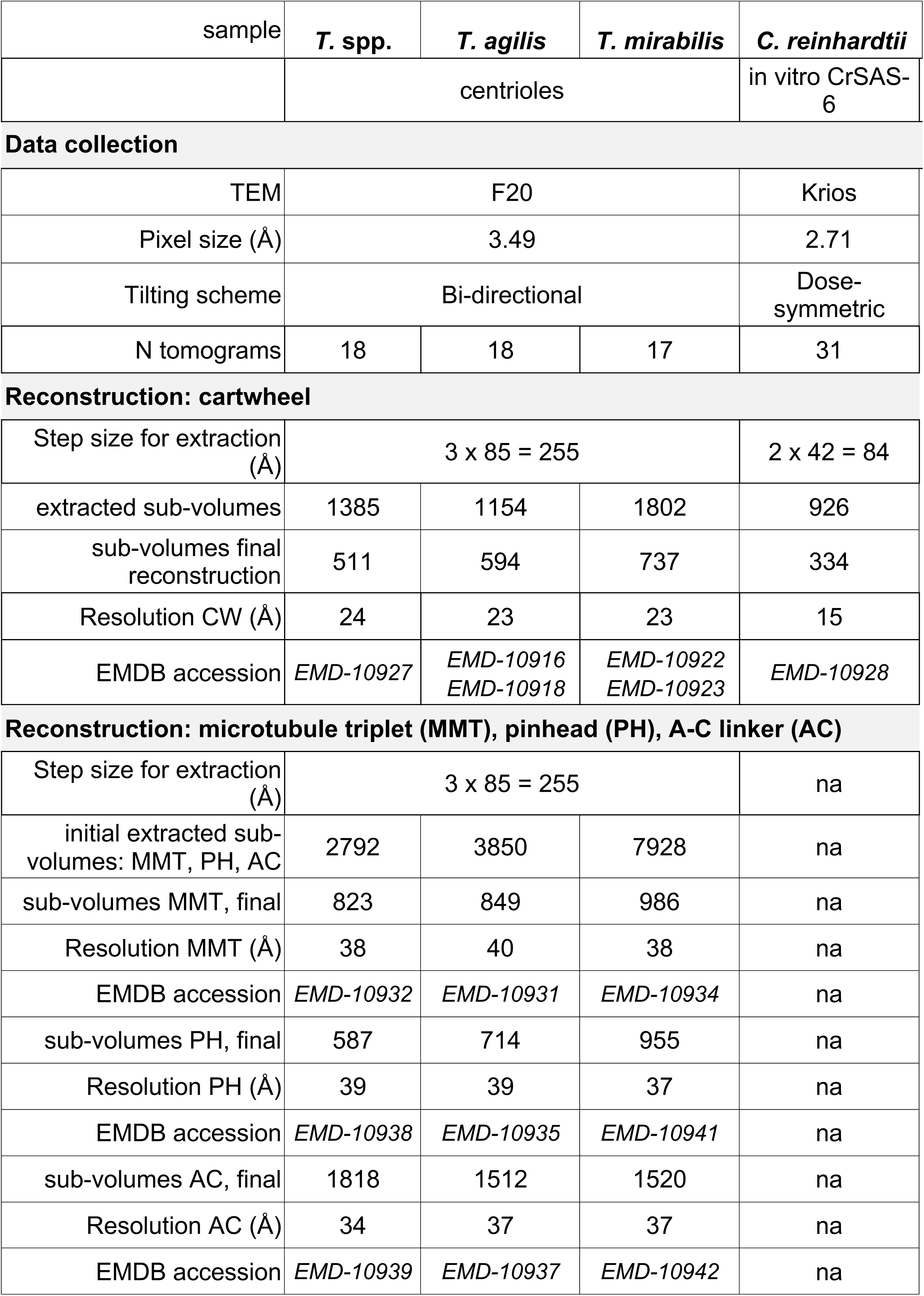
Statistics of Cryo-EM data collection and reconstruction and EMDB accession, see also Fig. S1.

**Table S2.** Correlation and overlap score for fitting with computationally assembled CrSAS-6[6HR] single, double, and double rings with offset. Results of the fit in map function performed with ChimeraX by maximizing correlation as shown in Fig. 5 and Fig. S9, Fig. S10 and described in Materials and Methods. Reported are the percentage of atoms inside the density map, the correlation, and the overlap between model and map. For each STA map, we report the gain in overlap upon fitting the double and offset double versus single ring model; the maximum possible gain is +100 %.

| Map | Ring model | Percentage (%) of atoms inside contour | Correlation | Overlap | Percentage (%) gain in overlap score by fitting double over single ring |
| --- | --- | --- | --- | --- | --- |
| ***T. mirabilis*** 36% | Single | 77 | 0.83 | 23 |  |
|  | Double | 63 | 0.71 | 31 | +58 |
|  | Offset | 69 | 0.73 | 27 | +87 |
| ***T. mirabilis*** 64% | Single | 93 | 0.78 | 17 |  |
|  | Double | 78 | 0.71 | 26 | +47 |
|  | Offset | 71 | 0.74 | 34 | +47 |
| ***T. spp.*** | Single | 94 | 0.83 | 34 |  |
|  | Double | 64 | 0.65 | 41 | +23 |
|  | Offset | 63 | 0.69 | 45 | +33 |
| ***T. agilis*** 55% | Single | 99 | 0.86 | 38 |  |
|  | Double | 82 | 0.73 | 51 | +34 |
|  | Offset | 86 | 0.78 | 55 | +44 |
| ***T. agilis*** 45% (thick) | Single | 100 | 0.86 | 41 |  |
|  | Double | 87 | 0.79 | 60 | +46 |
|  | Offset | 78 | 0.83 | 64 | +56 |
| ***T. agilis*** 45% (thin) | Single | 93 | 0.87 | 33 |  |
|  | Double | 46 | 0.63 | 34 | +4 |
|  | Offset | 46 | 0.63 | 34 | +4 |
| **CrSAS-6[NL]** in vitro | Single | 91 | 0.90 | 18 |  |
|  | Double | 91 | 0.90 | 35 | +97 |
|  | Offset | 84 | 0.88 | 33 | +83 |
| Correlation and overlap are calculated with ChimeraX (Goddard *et al.*, 2018).<br>Correlation value range: -1 to 1 |  |  |  |  |  |

### Supplemental Figure Legends

**Figure S1:**
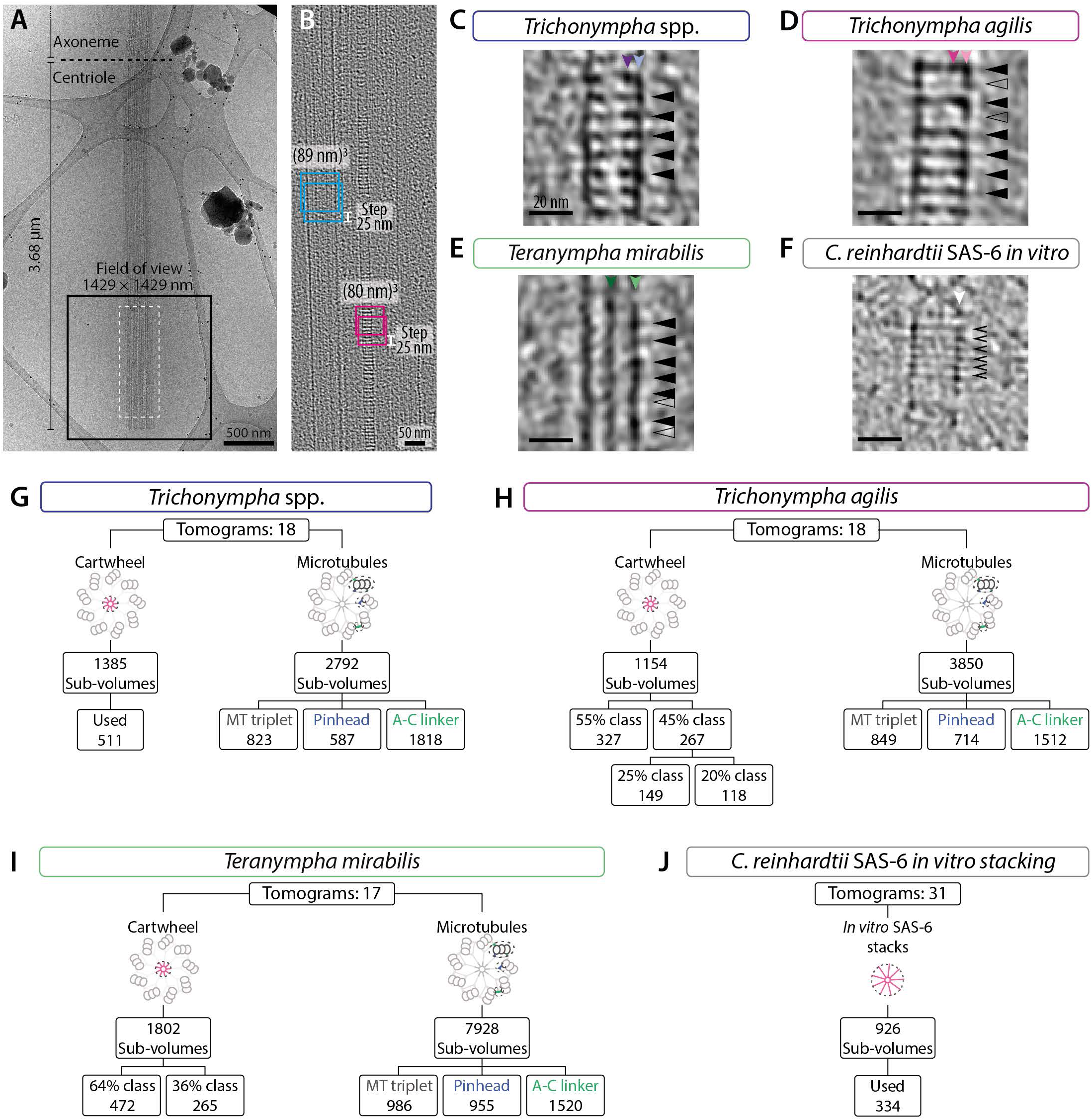
Cryo-EM acquisition, raw data and processing. (A) Low magnification view of cryo-preserved purified *T.* spp. centriole and axoneme on carbon grid. Note that the proximal end of the centriole is blunt, and that the diameter of the centriole containing microtubule triplets is larger than that of the axoneme containing microtubule doublets, enabling one to determine the length of the centriole, as indicated. The black square illustrates the proximal location and the 1429 × 1429 nm size of the field of view typically used for tilt series acquisition; note that this can be less than the length of the centriole depending on the species. Dashed white rectangle: area corresponding to the tomogram shown in panel B. (B) 2D longitudinal view (sum of 10-20 slices) of the central cartwheel tomogram, with band-pass filter and Gaussian blur corresponding to the area with a dashed rectangle in (A). Sub-volumes extracted along the longitudinal axis are (80 nm)^3^ for the central cartwheel (pink squares represent one plane of the sub-volume) and (89 nm) ^3^ for the peripheral elements comprising microtubule triplet, pinhead and A-C linker (blue squares represent one plane of the sub-volume), with a step size of 25 nm in both cases. (C-F) 2D longitudinal views from raw reconstructed tomograms of the central cartwheel illustrating typical sub-volumes (sum of 10-20 slices of the central cartwheel tomogram, with band-pass filter and Gaussian blur) from *T.* spp. (C), *T. agilis* (D), *T. mirabilis* (E) and CrSAS-6[NL] stacked assemblies (F). Filled arrowhead indicate hub elements constituted of double units that can sometimes be resolved upon STA, but are not visible in this raw data (C-E), empty arrowhead indicate hub elements in the *T. agilis* 45 % class lacking the CID (D). Note that classes cannot be unambiguously distinguished in the *T. mirabilis* raw tomograms, empty arrowheads in this case indicate hub elements with apparent variations in spacing (E). Chevrons indicate resolved CrSAS-6[NL] single unit hub elements (F). (G-J) Processing scheme with number of tomograms, initial sub-volumes extracted for cartwheel and peripheral elements (microtubule triplet, pinhead and A-C linker), classification/refinement steps, and final sub-volumes used for STA in *T.* spp. (G), *T. agilis* (H), *T. mirabilis* (I) and CrSAS-6[NL] stacked assemblies (J). See also Table S1.

**Figure S2:**
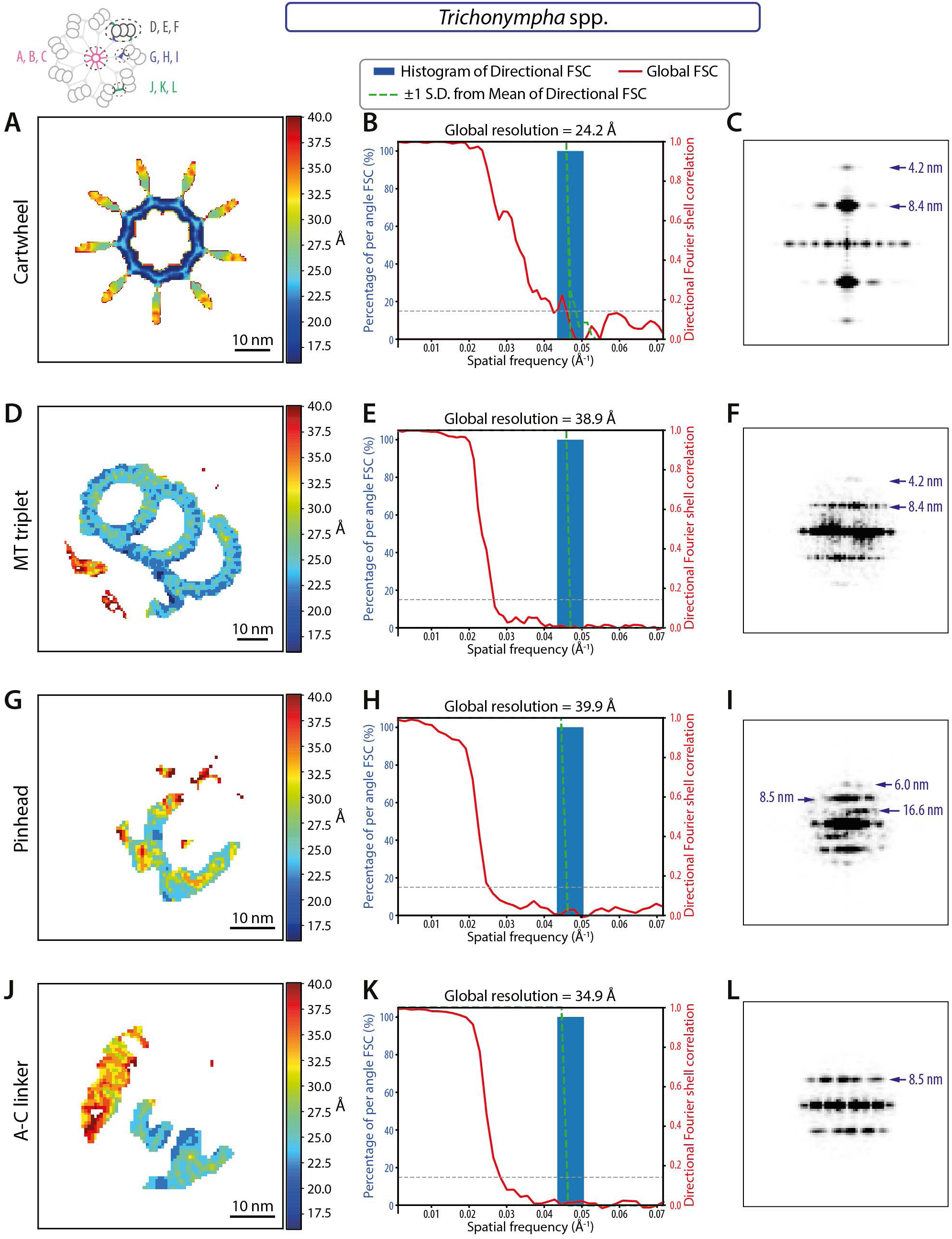
Resolution maps of *T.* spp. (A, D, G, J) Local resolution distribution in *T.* spp. STA estimated with ResMap. Shown are transverse views of central cartwheel (A), microtubule triplet (D), pinhead (G) and A-C linker (J). (B, E, H, K) Histogram and directional Fourier shell correlation (FSC) plots for *T.* spp. STA of central cartwheel (B), microtubule triplet (E), pinhead (H) and A-C linker (K) with global resolution at FSC 0.143 criterion indicated. (C, F, I, L) Power spectra of 2D class average with periodicities highlighted by arrows for *T.* spp. STA of central cartwheel (C), microtubule triplet (F), pinhead (I) and A-C linker (L). The precision for the measurements in this and remaining power spectra is 1 pixel, corresponding to a minimum of 0.4 nm (see Table S1).

**Figure S3:**
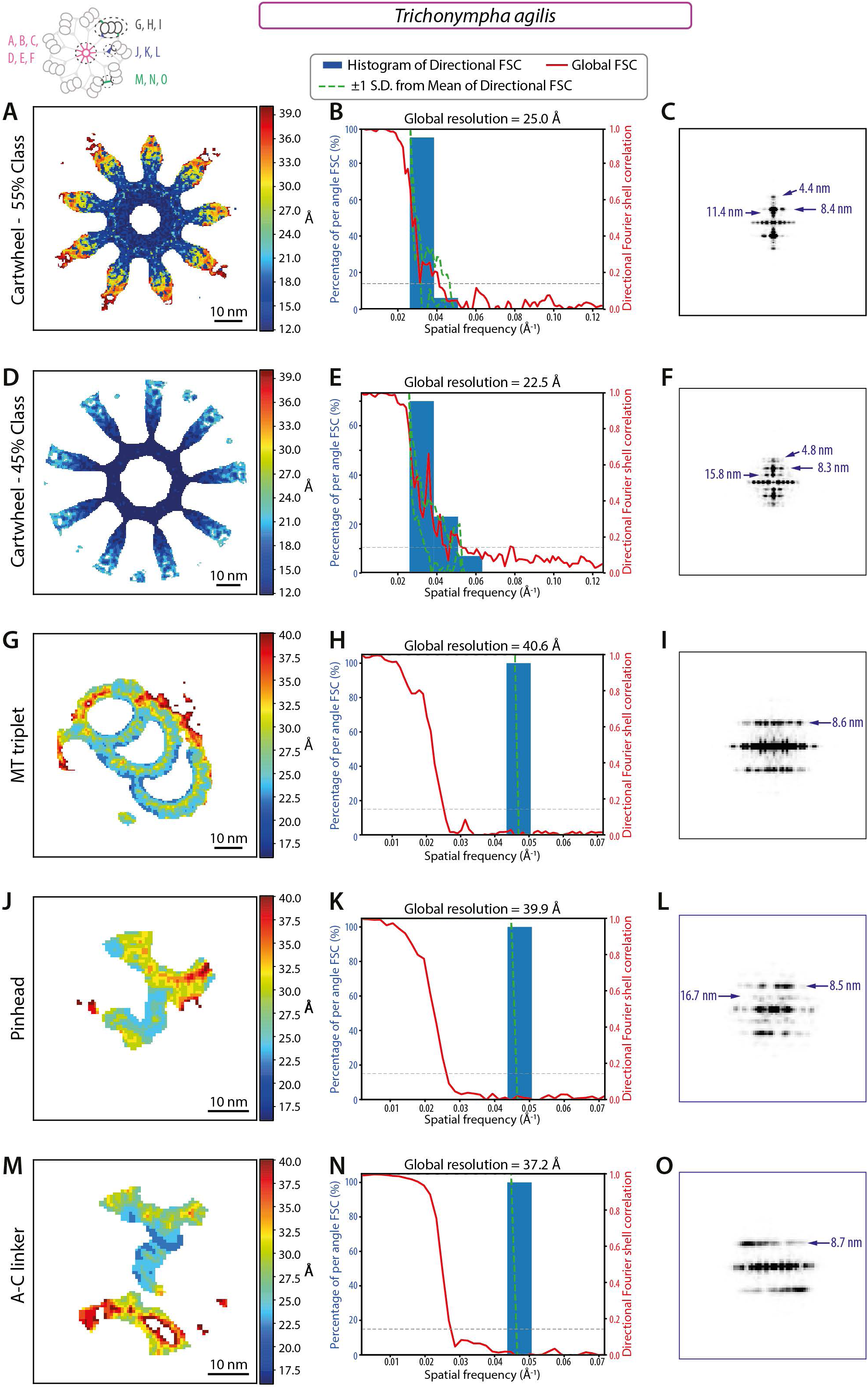
Resolution maps of *T. agilis*. (A, D, G, J, M) Local resolution distribution in *T. agilis* STA estimated with ResMap. Shown are transverse views of central cartwheel 55 % class (A), 45 % class (D), microtubule triplet (G), pinhead (J) and A-C linker (M). (B, E, H, K, N) Histogram and directional FSC plots for *T. agilis* STA of central cartwheel 55 % class (B), 45 % class (E), microtubule triplet (H), pinhead (K) and A-C linker (N) with global resolution at FSC 0.143 criterion indicated. (C, F, I, L, O) Power spectra of 2D class average with periodicities highlighted by arrows for *T. agilis* STA of central cartwheel 55 % class (C), 45 % class (F), microtubule triplet (I), pinhead (L) and A-C linker (O).

**Figure S4:**
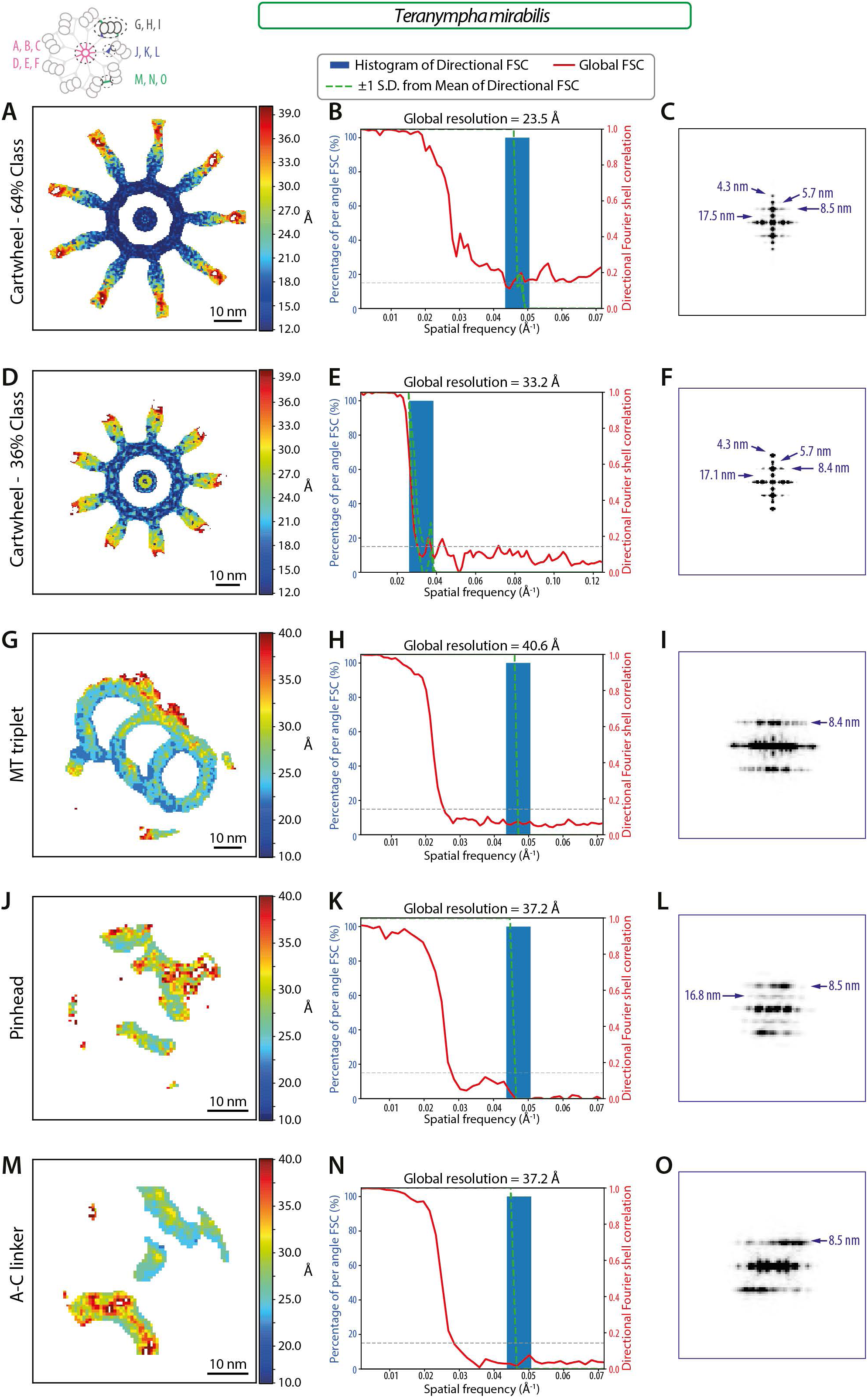
Resolution maps of *T. mirabilis*. (A, D, G, J, M) Local resolution distribution in *T. mirabilis* STA estimated with ResMap. Shown are transverse views of central cartwheel 64 % class (A), 36 % class (D), microtubule triplet (G), pinhead (J) and A-C linker (M). (B, E, H, K, N) Histogram and directional FSC plots for *T. mirabilis* STA of central cartwheel 64 % class (B), 36 % class (E), microtubule triplet (H), pinhead (K) and A-C linker (N) with global resolution at FSC 0.143 criterion indicated. (C, F, I, L, O) Power spectra of 2D class average with periodicities highlighted by arrows for *T. mirabilis* STA of central cartwheel 64 % class (C), 36 % class (F), microtubule triplet (I), pinhead (L) and A-C linker (O).

**Figure S5:**
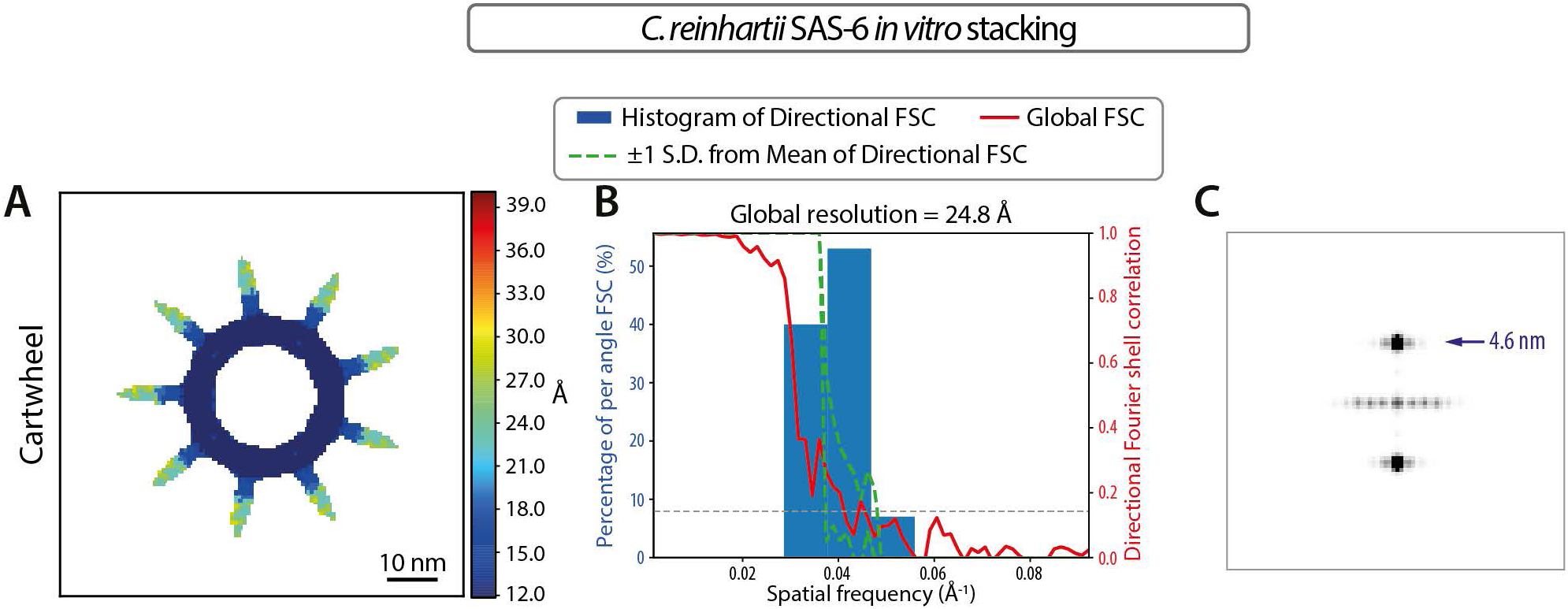
Resolution maps of CrSAS-6[NL] stacked assemblies. (A) Local resolution distribution in CrSAS-6[NL] stacked assemblies (generated from side-views) STA estimated with ResMap shown in transverse views. (B) Histogram and directional FSC plots for CrSAS-6[NL] stacked assemblies STA with global resolution at FSC 0.143 criterion indicated. (C) Power spectra of 2D class average with periodicity highlighted by arrows for CrSAS-6[NL] stacked assemblies.

**Figure S6:**
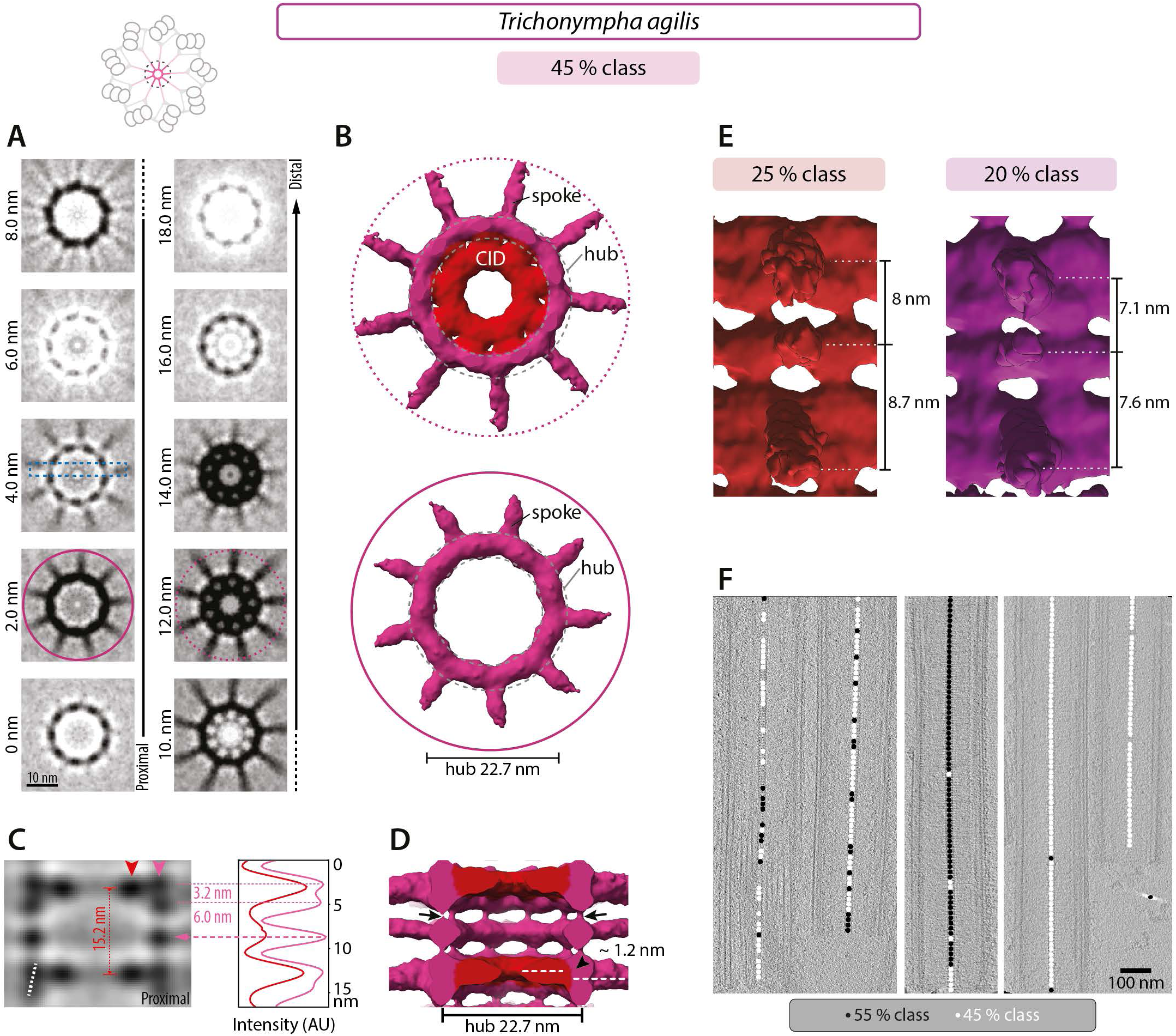
Variation in hub architecture is some *T. agilis* sub-volumes. (A) Transverse 2D slices through central cartwheel STA of *T. agilis* 45% class, which comprises 25 % and 20 % sub-classes (see E), at indicated height, from proximal (0 nm) to distal (18.0 nm). The pink circles mark spokes with CID (12.0 nm, dashed line) or without CID (2.0 nm, solid line), as represented in (B); dashed box in the 4.0 nm slice indicates longitudinal section shown in (C). Schematic on top illustrates the area used to generate the 3D maps of the central cartwheel. (B) Transverse views of central cartwheel STA 3D map of *T. agilis* 45 % class. The hub diameter is 22.7 nm ± 0.2 nm (N=3) with 9 emanating spoke densities, either at a level where the CID is present (dashed circle) or not (solid circle), as indicated in (A). (C) 2D longitudinal view of central cartwheel STA of *T. agilis* 45% class delineated by a dashed box in (A). Arrowheads denote position of line scans along the vertical axis at the level of the CID (dark pink) and the hub (light pink), with corresponding normalized pixel intensities in arbitrary units (AU). The plot profiles are shifted horizontally relative to each other for better visibility. The distance between hub densities alternates between 3.2 nm (N=1) and 6.0 ± 0.3 nm (N=2); maxima are indicated by dashed pink lines. Dashed white line indicates offset of two superimposed hub units. The average distance between two CID elements is 15.2 nm (N=1; dashed red line). The middle hub density that comprises only one unit and lacks a neighboring CID is indicated by a dashed arrow. (D) Longitudinal view of central cartwheel STA of *T. agilis* 45 % class at lower contour level than in (B). Note densities bridging successive hubs vertically (arrows), as well as proximal location of CID relative to the spoke density axis, resulting in a vertical offset of 1.2 ± 0 nm (N=2; arrowhead). Note also absence of CID in middle hub element comprising a single unit. (E) The spacing between the spoke density emanating from a double hub unit and the spoke density emanating from a single hub unit distal to it is 8.0 nm in the 25 % class and 7.1 nm in the 20 % class (both N=1). By contrast, the spacing between the spoke density emanating from a double hub unit and the spoke density emanating from a single hub unit proximal to it is 8.7 nm in the 25 % class and 7.6 nm in the 20 % class (both N=1). (F) Distribution of sub-volumes of the 55 % (black circle) and 45 % (white circle) classes along five *T. agilis* centrioles; areas with neither black nor white circle could not be clearly assigned to either class. Note individual centrioles constituted of mostly one of the two classes, and others where the two classes are mixed without an apparent pattern, indicating that the distribution of 55 % and 45 % classes is not stereotyped along the centriole.

**Figure S7:**
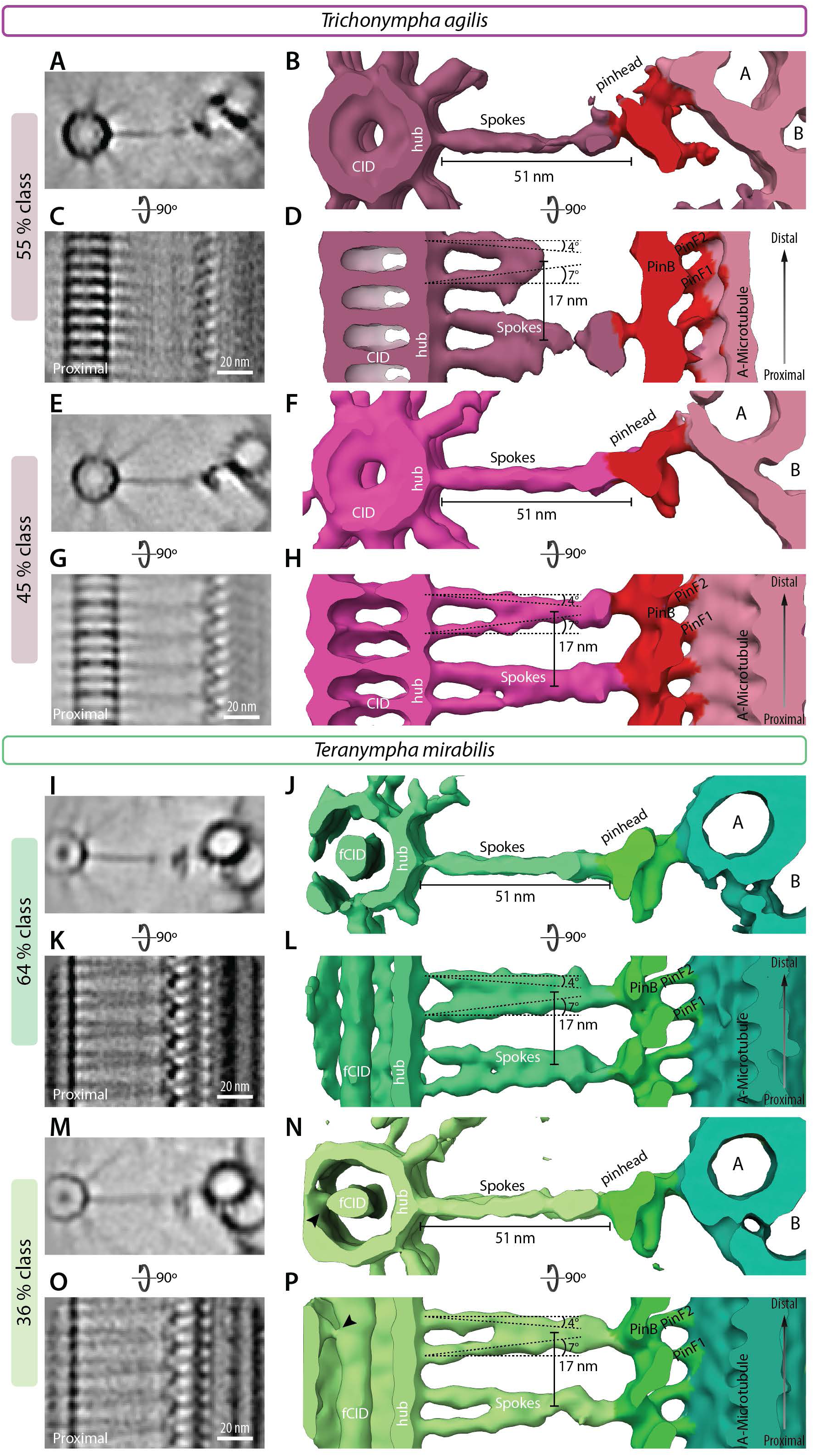
Polar centriolar cartwheel. (A-P) Non-symmetrized STA comprising larger volume than in Fig. 2-4 and centered on the spokes to jointly show the central cartwheel and peripheral elements in *T. agilis* (A-D: 55 % class; E-H: 45 % class) and *T. mirabilis* (I-L: 64 % class; M-P: 36 % class). 2D slices through STA transverse view (A, E, I, M) with corresponding 3D views (B, F, J, N), as well as 2D longitudinal views (C, G, K, O) with corresponding 3D views (D, H, L, P). The concerted proximal-distal polarity is visible from the central CID (in A-H) and the hub all the way to the pinhead. Note that proximal-distal polarity is visible also in the asymmetric spoke densities tilt angles in all cases, with a more pronounced tilt on the proximal side (D, H, L, P). Note also that the unsymmetrized CID shown here resembles that in the symmetrized maps of Figure 2, indicating that the CID exhibits *bona fide* 9-fold radial symmetry (A, E). Arrowhead in (N, P) points to a connection between fCID and hub. For representation, a Gaussian filter was applied to maps in ChimeraX.

**Figure S8:**
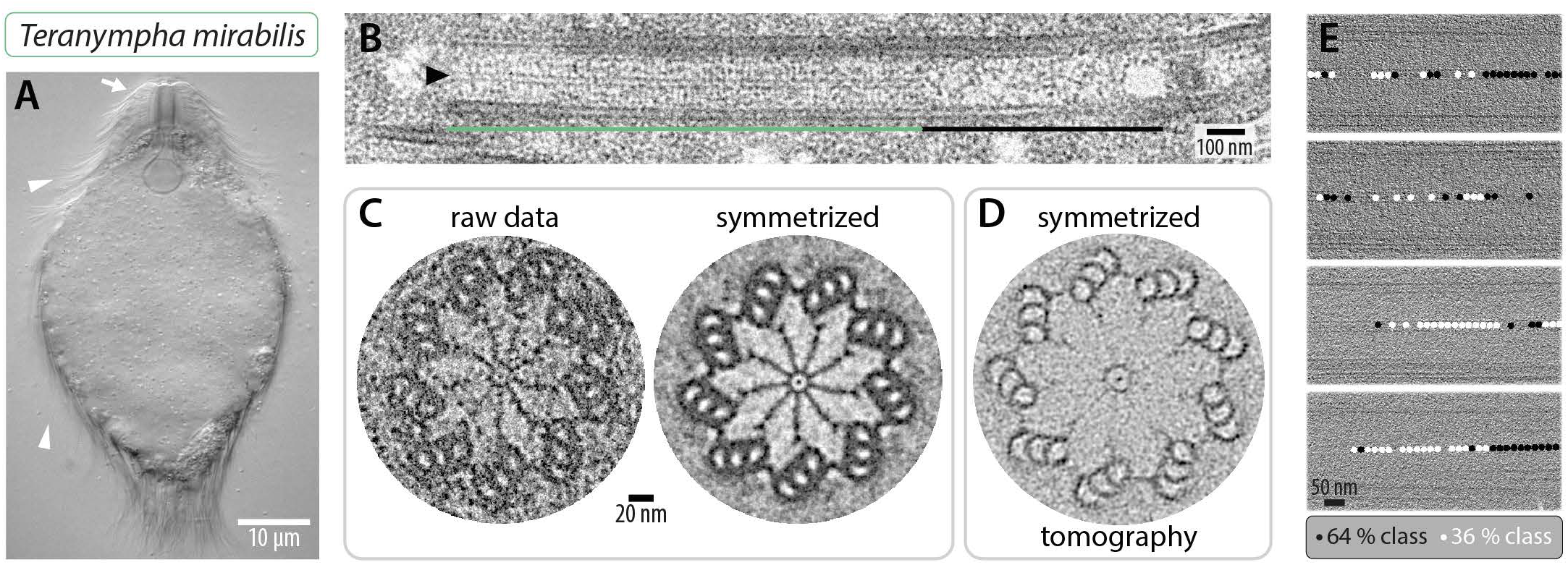
Very long cartwheel in *T. mirabilis.* (A) Differential interference contrast micrograph of live *T. mirabilis* cell. The arrow points to cell anterior, where the rostrum is located; arrowheads point to some of the flagella. (B) Transmission electron micrograph of *T. mirabilis* centriole embedded in resin – longitudinal view. The hub (arrowhead) is visible in the proximal cartwheel-bearing region (green line), but not in the distal region (black line). (C) Transmission electron micrograph of *T. mirabilis* centriole embedded in resin in transverse view (left) and corresponding image circularized and symmetrized (right). Note small density present inside the hub corresponding to the fCID. (D) Transverse view slice through cryo-electron tomogram of *T. mirabilis* centriole circularized and symmetrized. (E) Distribution of sub-volumes of the 64 % (black circle) and 36 % (white circle) classes along four *T. mirabilis* centrioles, proximal is left; areas with neither black nor white circle could not be clearly assigned to either class. Note that the distribution of the two classes is not stereotyped along the centriole.

**Figure S9:**
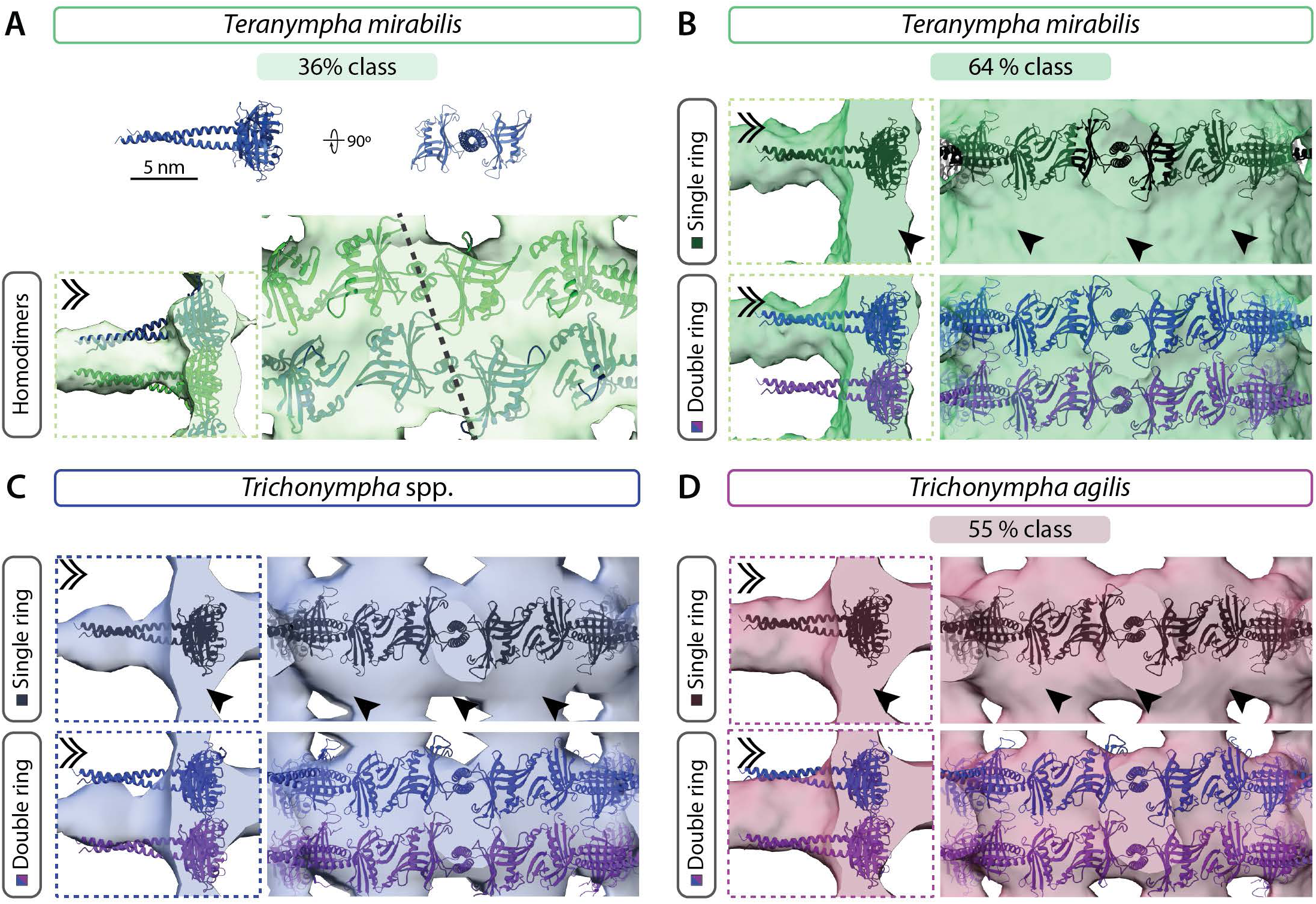
CrSAS-6[6HR] fitting in different cartwheel STA maps. (A) Ribbon representation of CrSAS-6[6HR] homodimers (top, scale bar 5nm) used for rigid body fitting into the 3D map from *T. mirabilis* 36 % class shown in side view (left) and end-on view magnified, with dashed line indicating offset between two superimposed homodimers (right). Individual layers of homodimers are indicated in distinct colors for clarity. In this and all other panels of this figure, dashed box indicates longitudinal section through hub element (left), double chevron indicates viewing direction of longitudinal external views (right). (B-D) CrSAS-6[6HR] single or double rings in register (ribbon diagram represented in different shades for clarity) fitted into the 3D maps from the *T. mirabilis* 64 % class (B), *T.* spp. (C) and the *T. agilis* 55 % class (D). Single ring fitting is shown in the top, double ring fitting at the bottom. Note unaccounted densities in the hub upon fitting of single rings in all cryo-ET maps (arrowheads, B-D), whereas all maps readily accommodate double CrSAS-6[6HR] rings at the hub element, although the coiled-coil elements extend slightly beyond the spoke densities. Note also that one coiled-coil in the *T. mirabilis* 64 % class sticks out of the density entirely (B, bottom).

**Figure S10:**
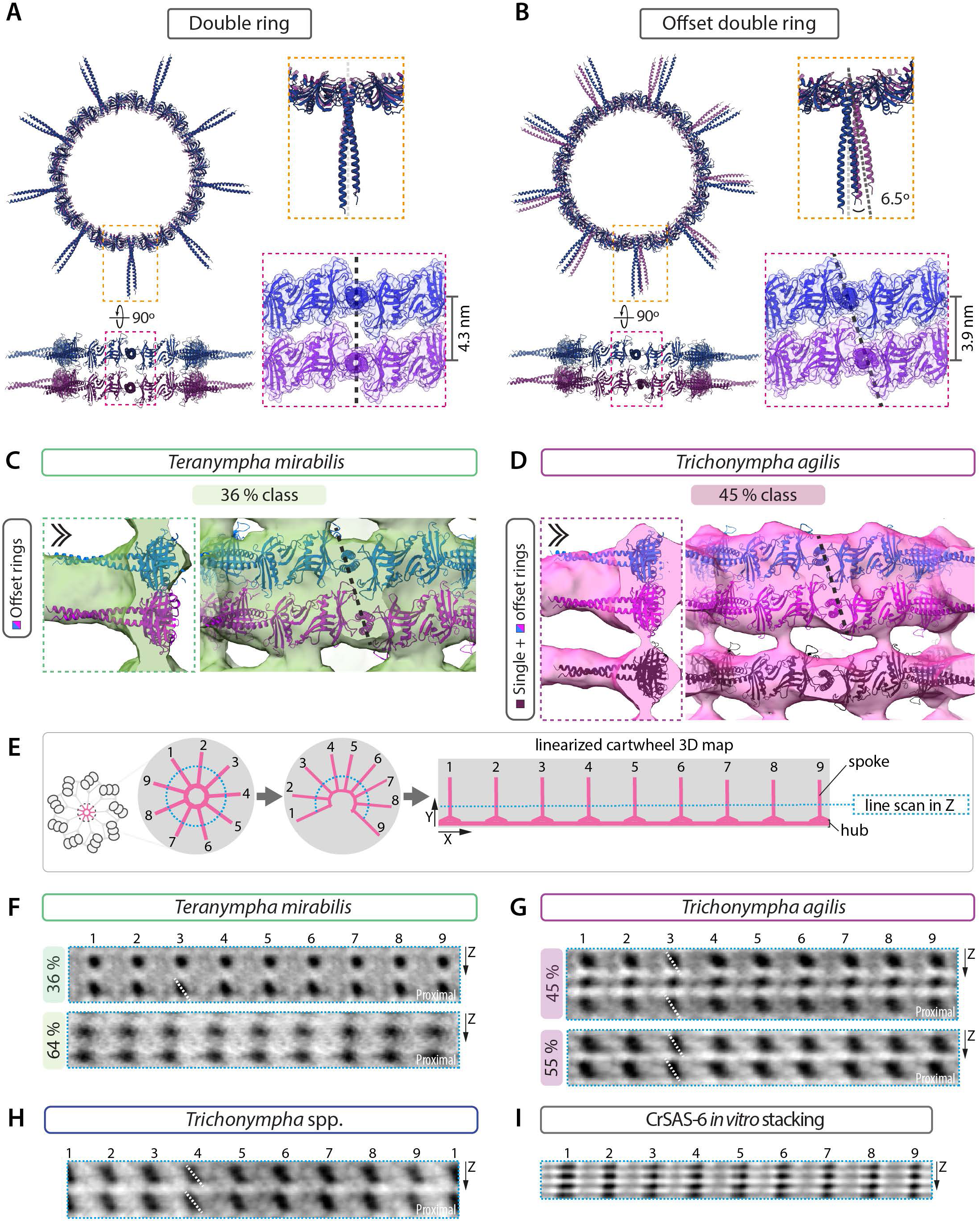
Superimposed rings of SAS-6 may be offset with respect to one another. (A, B) (left) Ribbon representation of CrSAS-6[6HR] homodimers computationally assembled into double rings (rings represented in different shades for clarity) with spokes in register (A) or with a 6.5° offset turning the proximal ring clockwise and moved 0.4 nm closer (B); transverse (top) and longitudinal view (bottom). (Right) Magnified views of two vertically superimposed homodimers (ribbon diagram), with dashed lines indicating angle between axes of the two coiled-coil axes (top) and surface representation of complementary interface (bottom). Distances were measured on the ribbon diagram. (C, D) CrSAS-6[6HR] offset double rings (shown in B and represented in different colors for clarity) fitted into 3D map from *T. mirabilis* 36 % class (C) and *T. agilis* 45% class (D), with dashed line indicating offset between double rings. Dashed box indicates longitudinal section through hub element (left), double chevron viewing point of longitudinal external views (right). (E) Schematic illustrating processing reported in (F-H): after linearization of the cartwheel STA, a longitudinal section in Z-direction (blue line) is shown at the level of the spokes (represented in pink). (F-H) Longitudinal section as explained in (E) for *T. mirabilis* 36 % and 64% classes (F), *T. agilis* 45 % and 55 % classes (G) and *T.* spp. (H) at the level of spoke emergence from the hub. Note offset of individual spokes highlighted by a dashed line. (I) Longitudinal section as explained in (E) for *in vitro* assembled CrSAS-6[NL] stacks; spokes from consecutive stacks are almost in register. Note that in (F-H) the spokes of double hub elements cannot be fully resolved, while they appear as individual units in (I).

